# In vivo reprogramming of cytotoxic effector CD8^+^ T cells via fractalkine-conjugated mRNA-LNP

**DOI:** 10.1101/2025.10.29.685358

**Authors:** Angela R. Corrigan, Shin Foong Ngiow, Maura Statzu, Maria Betina Pampena, Jayme M.L. Nordin, Amie Albertus, Stephen D. Carro, Justin Harper, Rachelle L. Stammen, Jennifer Wood, Jacob T. Hamilton, Houping Ni, Justin Su, Rajesvaran Ramalingam, Vincent H. Wu, Mirko Paiardini, Drew Weissman, E. John Wherry, Edward F. Kreider, Michael R. Betts

## Abstract

Selective in vivo reprogramming of cytotoxic effector CD8^+^ T (T_eff_) cells holds tremendous promise as a therapeutic tool but has not yet been accomplished. Here, we demonstrate that fractalkine-conjugated mRNA lipid nanoparticles (mRNA-LNP) can specifically target and deliver mRNA to CX3CR1^+^ T_eff_ cells in vitro and in vivo. In mice, fractalkine-conjugated LNP target up to 90% of blood and splenic T_eff_ cells, and delivery of IL-2-encoding mRNA to T_eff_ cells enables robust exogenous IL-2 secretion. In rhesus macaques, fractalkine-conjugated mRNA-LNP target up to ∼100% of peripheral blood T_eff_ cells and delivery of CD62L-mRNA enables transient CD62L expression. Collectively, these data demonstrate the potential of natural receptor ligand-based targeting of mRNA-LNP for effective and efficient transient in vivo modification of T_eff_ cells.

---

Cytotoxic effector CD8^+^ T (T_eff_) cells are a critical component of the adaptive immune response given their ability to kill virally infected or cancerous cells. As such, substantial efforts have been made to harness T_eff_ cells for a wide range of therapeutic purposes, such as cancer immunotherapy, autoimmunity, and chronic viral infection clearance (*1–4*). Recent clinical success of engineered chimeric antigen receptor T cells (CAR-T) highlights the ability of T cells to be successfully edited and/or reprogrammed for enhanced or altered immunological function (*5–7*). However, most lentiviral-based CAR-T approaches require ex vivo production and favor targeting and expansion of stem-like CD8^+^ T cell subsets, rather than T_eff_ cells (*8–10*). Importantly, CD8^+^ T cell differentiation to T_eff_ subsets is associated with repression of tissue homing receptors, including the requisite CD62L and CCR7 lymph node trafficking molecules (*11*), limiting the ability of peripheral blood T_eff_ cells to exit the vasculature and function in secondary lymphoid tissues, such as lymph nodes. Therefore, a robust method to directly target and modify intravascular T_eff_ cells in vivo and alter their trafficking patterns would broaden and revolutionize the therapeutic utility of T cell-based therapies. To date, however, there have not been any documented approaches to selectively target T_eff_ cells in vivo.

mRNA delivery via LNP is highly efficacious for expression of target proteins and represents a promising new approach for the transient in vivo modulation of T cells (*12–15*). However, mRNA-LNP are not readily taken up by lymphocytes (*16, 17*). To overcome this limitation, previous strategies to target T cells via mRNA-LNP have involved altering lipid formulations or conjugating T cell-specific monoclonal antibodies onto the surface of LNP, enabling their uptake through receptor-mediated endocytosis (*18–21*). While these strategies can broadly target CD4^+^ and/or CD8^+^ T cell populations, they lack specificity for distinct T cell subsets including T_eff_ cells. In addition, these antibody-conjugated mRNA-LNP have limited uptake and/or protein expression in target cell populations which may be attributed to suboptimal receptor mediated endocytosis upon antibody binding. As such, there remains a need to develop novel non-antibody-based methods to selectively target T_eff_ cells for in vivo reprogramming. As most intravascular T_eff_ cells can be distinguished from other CD8^+^ T cell subsets by the surface expression of CX3CR1/fractalkine receptor (*22, 23*), we hypothesized that the interaction between CX3CR1 and its cognate ligand, CX3CL1/fractalkine (*24, 25*), could be used to target CX3CR1^+^ T_eff_ cells with mRNA-LNP. Using this ligand-based mRNA-LNP approach, we then sought to reprogram T_eff_ cell tissue homing properties by delivering mRNA encoding the lymph node homing molecule CD62L to enable CX3CR1^+^ T_eff_ trafficking into secondary lymphoid tissues.

## Fractalkine-conjugated LNP targeting of T_eff_ cells in vitro

To determine whether fractalkine-conjugated mRNA-LNP could successfully target CX3CR1^+^ T_eff_ cells, we adapted previously described methods to conjugate recombinant soluble human fractalkine to green fluorescent protein (GFP)-encoding mRNA-LNP at different mRNA/fractalkine ratios (Fig 1A) (*18, 24, 26*). We then incubated the conjugated mRNA-LNP with purified peripheral human CD8^+^ T cells, total T cells (CD4^+^ and CD8^+^ T cells), or unfractionated peripheral blood mononuclear cells (PBMCs) and analyzed the expression of CX3CR1 and GFP in CD8^+^ T cells by flow cytometry across 20 different healthy donors (gating strategy shown in Fig S1A). After incubation with the fractalkine-conjugated LNP, CX3CR1 expression on the CD8^+^ T cells decreased concomitantly with an increase in GFP signal, indicative of LNP uptake via receptor mediated endocytosis and successful mRNA translation (Fig 1B). GFP expression was greatest in T_eff_ cells treated with LNP conjugated with the lowest tested fractalkine level (Fig1B-C and fig S1B-C). Using this 1:2 µg mRNA/fractalkine ratio, we found that the targeting of T_eff_ cells by these LNP and subsequent GFP expression was dose-dependent over a range of 0.125-1µg mRNA-LNP per million cells (Fig 1D). Similar results were also found when testing LNP with higher mRNA/fractalkine ratios (fig S1D-G). We observed rapid LNP uptake and mRNA translation, with GFP expression detected 16 hours post LNP addition (earliest timepoint analyzed) and with all T_eff_ cells expressing GFP within 40 hours (Fig 1E). Notably, the variation seen in the proportion of GFP^+^ CD8^+^ T cells across donors was significantly correlated to the level of CX3CR1 expression on the corresponding untreated CD8^+^ T cells (fig S1H). Collectively, these data indicate the successful targeting of human CX3CR1^+^ T_eff_ cells via fractalkine-conjugated mRNA-LNP in vitro and demonstrate that the LNP fractalkine content, dose, and timing can be optimized for increased LNP uptake and protein expression.

**Fig. 1.**
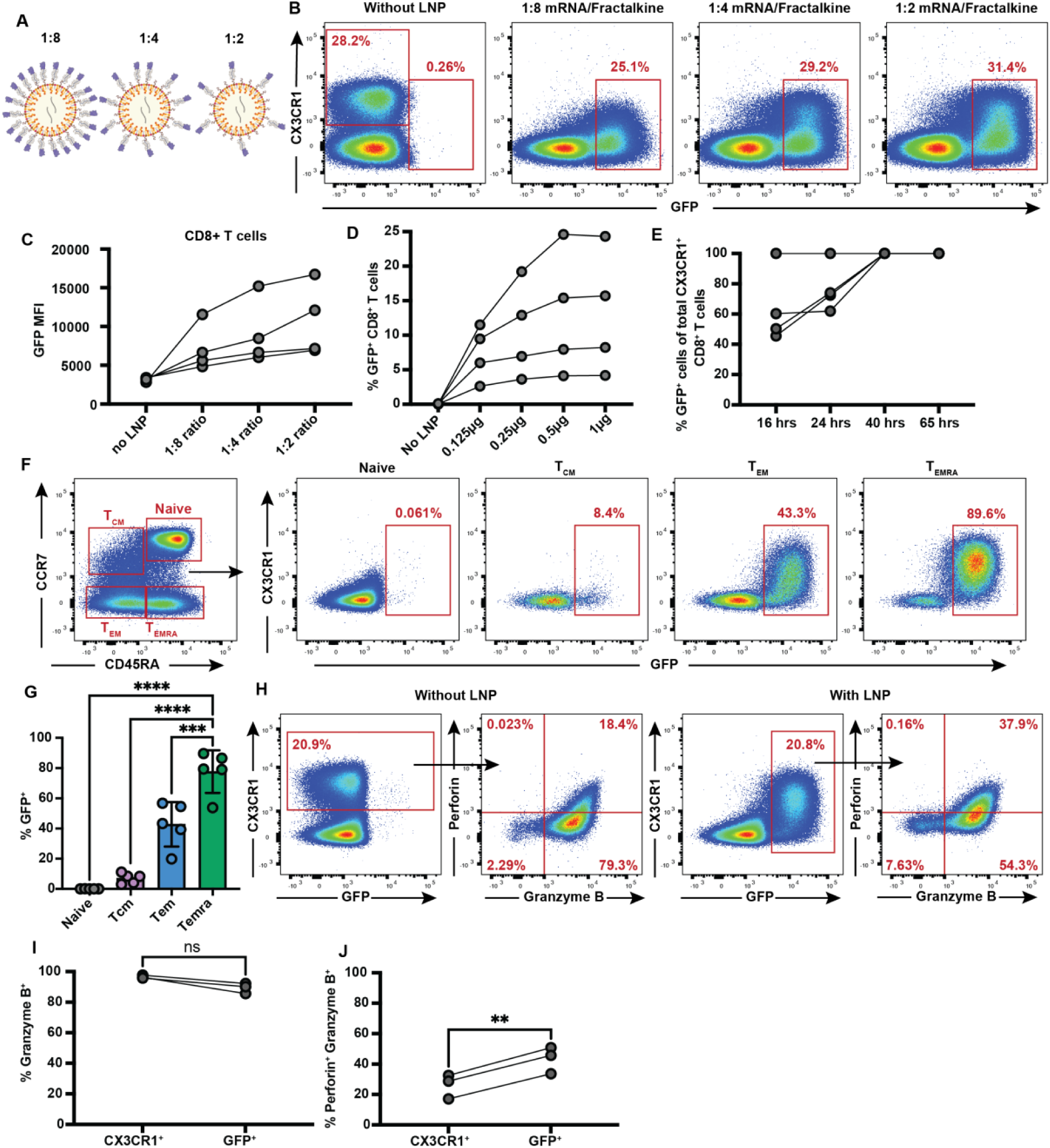
Fractalkine-conjugated LNP target CX3CR1^+^ T_eff_ cells in vitro: (A) Schematic representing recombinant human fractalkine-conjugated mRNA-LNP created with mRNA/ fractalkine ratios of 1:8, 1:4, and 1:2μg. (B) Flow cytometry analysis of human CD8^+^ T cells incubated for 65 hours with or without fractalkine-conjugated GFP-encoding mRNA-LNP shown in (A) at a dose of 0.5µg mRNA/million cells. Expression of CX3CR1 and GFP expression in the CD8^+^ T cells from one representative donor. (C) Median fluorescence intensity (MFI) of GFP from four donors tested. (D) Percentage of CD8^+^ T cells expressing GFP after incubation with or without fractalkine-conjugated GFP-encoding mRNA-LNP at 1:2 ratio at doses ranging from 0.125 to 1µg mRNA/million cells in four additional donors. (E) Percentage of CX3CR1^+^ T_eff_ cells expressing GFP in PBMC 16 to 65 hours post incubation with the 1:2 ratio fractalkine-conjugated GFP LNP in four additional donors. (F) Memory subset analysis of GFP expression patterns in canonical CD8^+^ T cell subsets, including, naïve (CCR7^+^/CD45RA^-^), central memory (T_CM_; CCR7^-^/CD45RA^-^), effector memory (T_EM_; CCR7^-^/CD45RA^+^) and terminally differentiated effectors (T_EMRA_; CCR7^+^/CD45RA^+^) after 65-hour incubation with fractalkine-conjugated GFP-encoding mRNA-LNP at 1:2 mRNA/Fractalkine ratio. (G) GFP expression in each of the four CD8^+^ T cell subsets from (F). (H) Perforin and granzyme B expression in CX3CR1^+^ CD8^+^ T cells incubated for 65 hours alone (left two panels) or in GFP^+^ CD8^+^ T cells incubated with fractalkine-conjugated GFP-encoding mRNA-LNP at a dose of 0.5µg mRNA/million cells (1:2 ratio) for 65 hours (right two panels). Data from one representative donor shown. (I) Percentage of cells expressing granzyme B or (J) both perforin and granzyme B for all donors tested. For all graphs, individual data points represent different donors. For all statistical analyses, *p < 0.05, **p < 0.01, ***p < 0.001, ****p < 0.0001; ns, not significant.

To evaluate the specificity of this platform to CX3CR1^+^ T_eff_ cell subsets, we evaluated GFP expression in naïve, central memory (T_CM_), effector memory (T_EM_), and CD45RA^+^ effector memory (T_EMRA_) CD8^+^ T cells. We observed GFP expression only in CD8^+^ populations known to express CX3CR1 (T_EM_, T_EMRA_ and a low proportion of T_CM_, but not naïve), with minimal off-target uptake by CX3CR1^-^ T cells (Fig 1F-G). Fractalkine-conjugated LNP-treated CD8^+^ T cells also had minimal disruption of perforin and granzyme B when compared to expression in untreated CX3CR1^+^ T_eff_ cells (Fig 1H-J), suggesting that LNP treatment does not induce T cell activation and degranulation. As CX3CR1 is also found on cytotoxic CD4^+^ T cells and CD56^dim^CD16^+^ cytotoxic natural killer (NK) cells (*24*), we assessed LNP uptake and mRNA translation in these cells (gating strategy shown in fig S1A and 3A, respectively). After fractalkine-conjugated GFP LNP treatment, we observed efficient targeting and mRNA delivery without any apparent change in the expression of granzyme B or perforin in these cell types (cytotoxic CD4^+^ T cells, fig S2A-D; CD16^+^ NK cells, S3B-E). We observed a similar distribution of GFP expression in CD4^+^ T cell subsets as seen in CD8^+^ T cells, such that the GFP expression was only highly detected in T_EM_ and T_EMRA_ CD4^+^ T cells (fig S2A-B). Within NK cell subsets, GFP expression was restricted to CD56^dim^CD16^+^ NK cells compared to CD56^bright^CD16^-^ regulatory NK cells further highlighting the specificity of fractalkine-conjugated LNP targeting to cells expressing CX3CR1 (*27*) (fig S3F-G).

Finally, to determine whether cellular signaling occurred downstream of fractalkine-conjugated LNP uptake, we performed single cell RNA sequencing on PBMCs that were either untreated, treated with unconjugated GFP LNP, or treated with fractalkine-conjugated GFP LNP (fig S4A). Besides GFP, we did not observe any significant differential gene expression relating to fractalkine/CX3CR1 signaling or cellular degranulation (fig S4B-C). This suggests that delivery of fractalkine-conjugated mRNA-LNP to CX3CR1^+^ cells does not activate signaling pathways downstream of CX3CR1 ligation or cytotoxicity pathways (fig S4D-E).

## Mouse fractalkine-conjugated LNP target CX3CR1^+^ T_eff_ cells in vivo

We next tested whether fractalkine-conjugated mRNA-LNP could target CX3CR1^+^ T_eff_ cells directly in vivo. As CX3CR1^+^ T_eff_ cells generally arise after antigen encounter, we used the lymphocytic choriomeningitis virus (LCMV)-Armstrong acute infection model in C57BL/6 mice, which induces significant expansion of circulating CX3CR1^+^ T_eff_ cells following infection (*22, 28*). Eight days post-LCMV infection, we administered 10μg of mouse fractalkine (chemokine domain only)-conjugated GFP LNP, unconjugated GFP LNP, or PBS intravenously (Fig 2A). After 24 hours, we analyzed GFP expression in peripheral blood and splenic CD8^+^ CX3CR1^+^ T_eff_ cells as well as H-2D^b^-GP_33-41_^+^ tetramer-specific (GP33^+^) CX3CR1^+^ T_eff_ cells (gating tree shown in fig S5). Compared to mice that received unconjugated LNP, we observed a ∼5 and 5.5-fold increase in peripheral blood GFP^+^ CX3CR1^+^ T_eff_ cells and GFP^+^ GP33^+^ CX3CR1^+^ T_eff_ cells, respectively (Fig 2B-C and fig S6A-B), in mice receiving fractalkine-conjugated LNP. We also observed a ∼2-fold increase in GFP expression within total splenic CX3CR1^+^ CD8^+^ and GP33^+^ CX3CR1^+^ T_eff_ cells over unconjugated LNP treatment, but we noted higher background GFP expression from the unconjugated LNP in spleen compared to peripheral blood (fig S6A-D). Additionally, fractalkine-conjugated LNP did not induce detectable CX3CR1 downregulation as observed for human cells in vitro (Fig 2B). In both PBMC and spleen, GFP expression in GP33^+^ CX3CR1^+^ T_eff_ cells was directly correlated with GFP expression in all CX3CR1^+^ CD8^+^ T cells. GFP expression in PBMC was also directly correlated with GFP expression in spleen (fig S6E-H). Furthermore, compared to unconjugated LNP, mice that received fractalkine-conjugated LNP had significantly higher GFP expression in all CX3CR1^+^ splenocytes, but not in CX3CR1^-^ splenocytes (fig S6I-K), indicating that fractalkine-conjugation does not alter nonspecific uptake of mRNA-LNP in CX3CR1^-^ splenocytes. Finally, LNP administration to mice in the memory phase after LCMV clearance (33 days post infection) also showed robust expression of GFP in GP33^+^ CX3CR1^+^ T_eff_ cells (Fig 2D and fig S7A-B), indicating that in vivo uptake of fractalkine-conjugated LNP in CX3CR1^+^ T_eff_ cells is not limited to acute infection or recent activation.

**Fig. 2.**
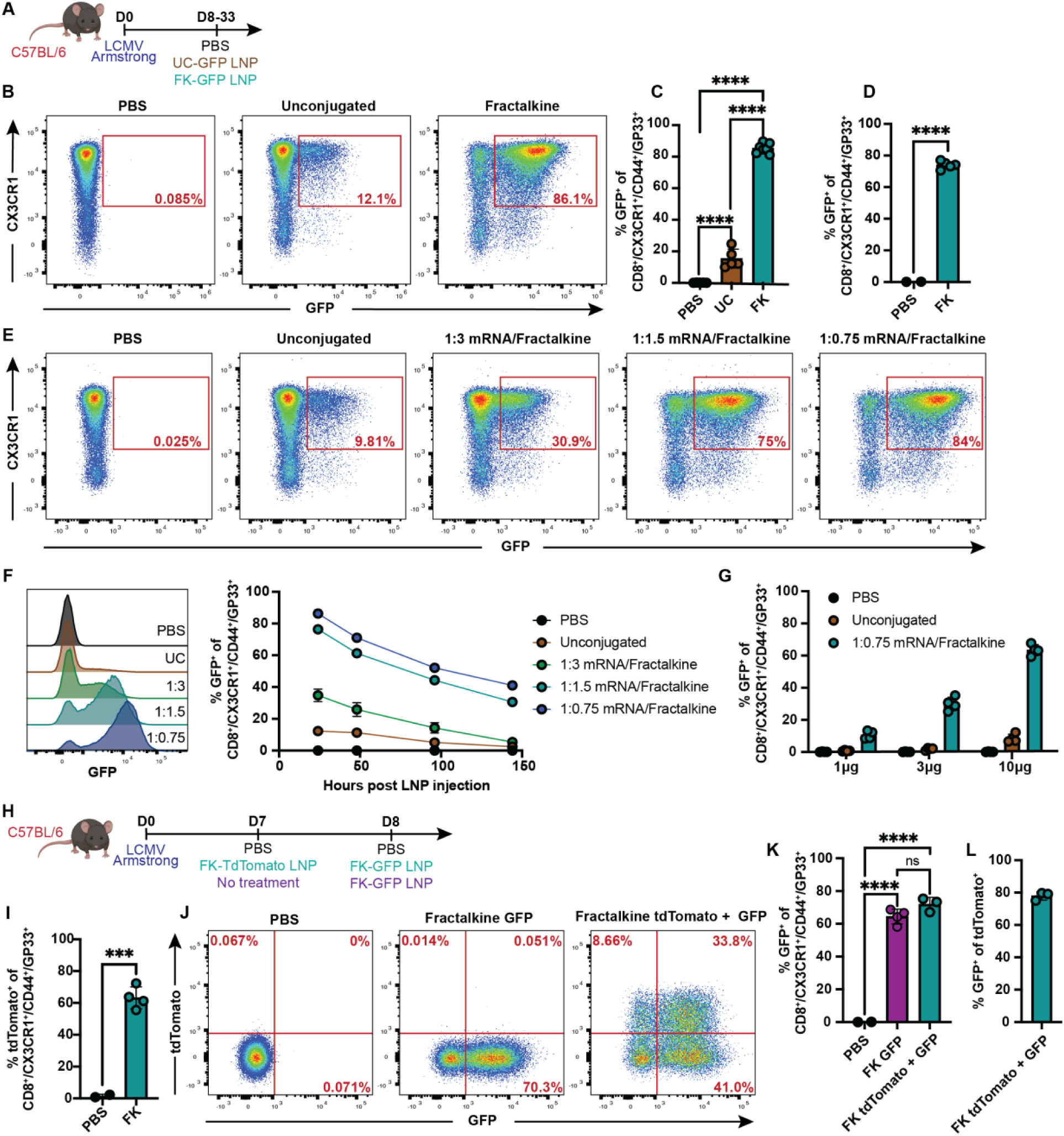
Mouse fractalkine-conjugated LNP target CX3CR1^+^ CD8^+^ T cells in vivo: (A) Experimental design for in vivo testing of mouse fractalkine-conjugated GFP-encoding mRNA-LNP. UC = unconjugated, FK = fractalkine-conjugated. (B) GFP expression in peripheral blood CX3CR1^+^/GP33^+^/CD44^+^/CD8^+^ T cells 24 hours after PBS or LNP injection (10μg dosage, 1:1.5 μg mRNA/μg fractalkine ratio, IV delivery); one mouse per group shown. Mice were infected with LCMV Armstrong for 8 days prior to treatment. (C) GFP expression from (B), quantified for all mice included in study. (D) Percentage of peripheral blood CX3CR1^+^/GP33^+^/CD44^+^/CD8^+^ T cells expressing GFP on day 34 post infection, 24 hours post LNP injection (10μg dosage, 1:0.75 ratio, IV delivery). (E) GFP expression in peripheral blood CX3CR1^+^/GP33^+^/CD44^+^/CD8^+^ T cells 24 hours after PBS or LNP injection (10μg dosage, IV delivery) with mouse fractalkine-conjugated GFP LNP made with 1:3, 1:1.5, or 1:0.75 μg mRNA/μg fractalkine ratio. Mice were infected with LCMV Armstrong for 8 days prior to treatment. Results from one mouse per group shown. (F) Representative histograms for GFP signal within T_eff_ cells from (E) at the 24-hour timepoint with one mouse per LNP treatment condition shown (left panel). Percentage of CX3CR1^+^/GP33^+^/CD44^+^/CD8^+^ T cells expressing GFP from (E) at each of the timepoints analyzed for all mice included in study. Mean ^+^ SD shown (right panel). (G) Percentage of peripheral blood CX3CR1^+^/GP33^+^/CD44^+^/CD8^+^ T cells expressing GFP 24 hours after injection with PBS, unconjugated or mouse fractalkine-conjugated GFP LNP at dosages of 1μg, 3μg, or 10μg (1:0.75 ratio, IV delivery). Mice were infected with LCMV Armstrong for 8 days prior to treatment. (H) Experimental design for data shown in (I-L) (10μg LNP dosage, 1:0.75 ratio, IV delivery). (I) Percentage of peripheral blood CX3CR1^+^/GP33^+^/CD44^+^/CD8^+^ T cells expressing tdTomato on day 8. (J) GFP and tdTomato expression in peripheral blood CX3CR1^+^/GP33^+^/CD44^+^/CD8^+^ T cells on day 9 post-infection. Results from one mouse per group shown. (K) GFP expression from (J) quantified for all mice included in study. (L) Percentage of peripheral blood tdTomato^+^/CX3CR1^+^/GP33^+^/CD44^+^/CD8^+^ T cells also expressing GFP on day 9 for all mice included in study. *p < 0.05, **p < 0.01, ***p < 0.001, ****p < 0.0001; ns, not significant. (B-D) represent one of two experimental repeats with similar results.

To further optimize in vivo mRNA delivery, we next tested different mRNA/fractalkine LNP conjugation ratios using the LCMV model and assessed GFP expression level and longevity. Similar to our in vitro findings, mice that received the fractalkine-conjugated LNP with the lowest mRNA/fractalkine ratio (1:0.75) had the highest proportion of GFP^+^ GP33^+^ CX3CR1^+^ T_eff_ cells at all timepoints analyzed, and GFP remained detectable six days post LNP injection at the lowest two conjugation ratios (Fig 2E-F and fig S8A-H). Using the lowest conjugation ratio, we then conducted a dose response experiment. On day eight post LCMV infection, we administered unconjugated or fractalkine-conjugated GFP mRNA-LNP at 1, 3, or 10μg dosages and examined GFP expression level in GP33^+^ CX3CR1^+^ T_eff_ cells after 24 hours. Compared to the 10μg dose, we found ∼2 and 6-fold decreases in GFP expression with the 3μg and 1μg dosages respectively (Fig 2G and fig S9A-B). However, we noted that the level of background uptake of unconjugated LNP in GP33^+^ CX3CR1^+^ T_eff_ cells relative to fractalkine-conjugated LNP decreased with smaller doses, leading to heightened specificity (fig S9C). Together, these data demonstrate that LNP fractalkine content and dose can be optimized for increased LNP uptake and protein expression in vivo.

Unlike our human in vitro targeting, intravenous administration of mouse fractalkine-conjugated LNP did not lead to downregulation of CX3CR1 in CD8^+^ T cells. This enabled us to assess whether we could sequentially deliver different fractalkine-conjugated mRNA-LNP to CX3CR1^+^ T_eff_ cells in vivo. Seven days post LCMV infection, mice were given PBS (group 1), fractalkine-conjugated tdTomato mRNA-LNP (group 2), or no treatment (group 3) (Fig 2H). The following day, we observed robust tdTomato expression in peripheral blood GP33^+^ CX3CR1^+^ T_eff_ cells after delivery of fractalkine-conjugated LNP but not PBS (Fig 2I and fig S10A). The mice then either received another dose of PBS (group 1) or fractalkine-conjugated GFP mRNA-LNP (groups 2-3) (Fig 2H). After 24 hours, we analyzed the expression of tdTomato and GFP in CX3CR1^+^ T_eff_ cells and found that GFP expression was similar between mice that received prior fractalkine-conjugated tdTomato LNP and mice that only received fractalkine-conjugated GFP LNP (Fig 2J-K and fig S10B). Additionally, approximately 78% of the tdTomato^+^ CX3CR1^+^ T_eff_ cells from peripheral blood and 85% of those from spleen expressed GFP, indicating that prior LNP administration did not interfere with sequential targeting of the same CX3CR1^+^ T_eff_ cells in vivo (Fig 2J, L and fig S10C).

## In vivo manipulation of mouse CX3CR1^+^ T_eff_ cells

We next examined whether fractalkine-conjugated mRNA-LNP could enable the production of a biologically active protein in vivo. As CX3CR1^+^ T_eff_ cells do not secrete IL-2 following clearance of LCMV infection (*29, 30*), we tested whether fractalkine-conjugated IL-2-encoding mRNA-LNP (fractalkine IL-2 mRNA-LNP) could drive exogenous IL-2 production in CX3CR1^+^ T_eff_ cells. Following viral clearance on day eleven post LCMV infection, we administered either fractalkine IL-2 mRNA-LNP, unconjugated IL-2 mRNA-LNP, or PBS (Fig 3A). Three hours post treatment, the mice were sacrificed, and serum IL-2 levels were analyzed via ELISA. Mice that received unconjugated LNP and fractalkine-conjugated LNP had high serum levels of IL-2 (Fig 3B). While IL-2 produced after delivery of unconjugated IL-2 LNP likely originated from ApoE-mediated uptake of LNP by hepatocytes (*16, 31*), it was unclear if CX3CR1^+^ T_eff_ cells contributed to the serum IL-2 after delivery of fractalkine IL-2 mRNA-LNP. We therefore sorted splenic CX3CR1^+^ and CX3CR1^-^ CD8^+^ T cells, cultured them for three hours in vitro, and compared IL-2 levels in the supernatant by ELISA. We found that only the CX3CR1^+^ CD8^+^ T cells from mice that received fractalkine-conjugated LNP produced IL-2 (Fig 3C). To directly demonstrate IL-2 production by splenic CX3CR1^+^ T_eff_ cells following LNP delivery, we assessed IL-2 production by flow cytometry. While IL-2 was largely undetectable in CX3CR1^+^ T_eff_ cells of mice that received PBS or unconjugated IL-2 LNP, ∼70% of CX3CR1^+^ T_eff_ cells produced IL-2 in mice that received the fractalkine-conjugated IL-2 LNP (Fig 3D-E). Collectively, these data highlight the ability of the fractalkine-conjugated LNP to both rapidly target and transiently reprogram CX3CR1^+^ T_eff_ cells in vivo.

**Fig. 3.**
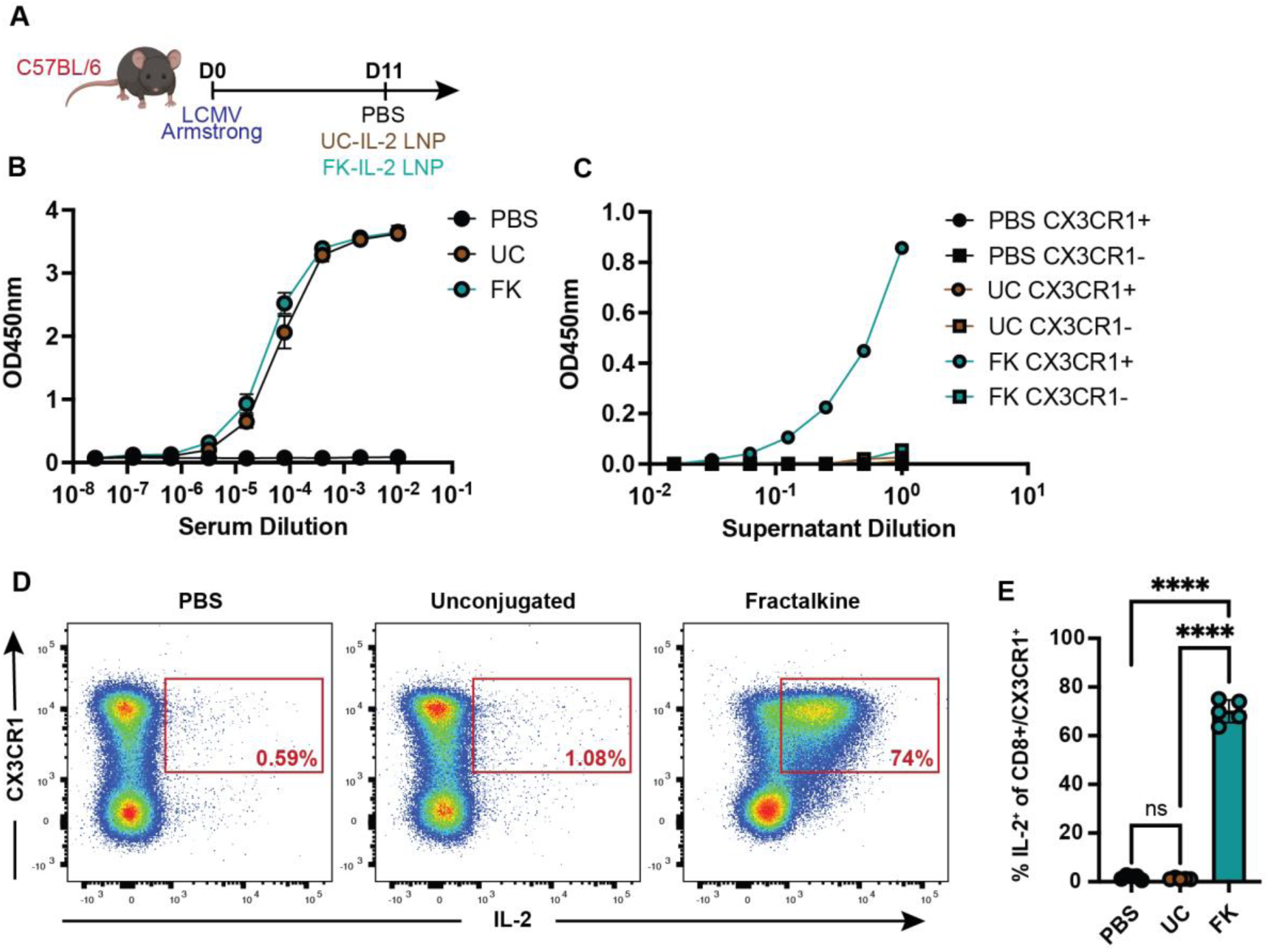
CX3CR1^+^ T_eff_ cells secrete IL-2 following fractalkine-conjugated IL-2 mRNA-LNP injection: (A) Experimental design for in vivo testing of mouse fractalkine-conjugated IL-2-encoding mRNA-LNP (6.5μg dosage, 1:1.5 μg mRNA/μg fractalkine ratio, IV delivery). UC = unconjugated, FK = fractalkine-conjugated. Mice were infected with LCMV Armstrong for 11 days prior to treatment and sacrificed three hours post LNP injection. (B) Serum IL-2 levels three hours post LNP injection tested by ELISA as measured by OD450nm plotted against serum dilution factors for all mice included in study. Mean with standard deviation for each group shown. Data show one of two replicative experiments. (C) Supernatant IL-2 levels from sorted splenic CX3CR1^+^ and CX3CR1^-^ CD8^+^ T cells after two hours, tested by ELISA as measured by OD450nm plotted against supernatant dilution factors. Splenocytes from all mice per group were combined prior to sorting. Data show one of two representative experiments. (D) Flow cytometry analysis of IL-2 secretion from CX3CR1^+^ CD8^+^ T cells three hours after PBS or LNP injection with results from one mouse per group shown. (E) Percentage of CX3CR1^+^ CD8^+^ T cells secreting IL-2 from (D), each symbol represents one mouse.

## Fractalkine-conjugated LNP target rhesus macaque CX3CR1^+^ T_eff_ cells in vivo

To assess the clinical translatability of this platform, we next tested the targeting of CX3CR1^+^ T_eff_ cells by human fractalkine-conjugated GFP mRNA-LNP in rhesus macaques (RMs, Fig 4A). Two RMs received a single dose of human fractalkine-conjugated GFP-encoding mRNA-LNP intravenously, and we monitored GFP expression in CX3CR1^+^ T_eff_ cells longitudinally (Fig 4B-C and fig S11A). One day post infusion, we observed a complete loss of peripheral blood CD8^+^ T cell CX3CR1 expression in both animals, concomitant with a reciprocal increase in GFP expression, suggesting ∼100% targeting of circulating CX3CR1^+^ T_eff_ cells. GFP remained detectable in CD8^+^ T cells for up to 14 days, with progressive recovery of CX3CR1 expression (Fig 4B-D). We further analyzed GFP expression within naïve, T_CM_, and T_EM_ CD8^+^ T cell populations and found GFP expression was largely restricted to the T_EM_ subset, indicating high specificity of fractalkine-conjugated LNP to rhesus CD8^+^ T cells expressing CX3CR1 (Fig 4E-G). Similar to CD8^+^ T cells, the fractalkine-conjugated LNP also targeted ∼100% of circulating CX3CR1^+^ NK cells as demonstrated by a complete loss of CX3CR1 staining and reciprocal increase in GFP expression (fig S11A-D).

**Fig. 4.**
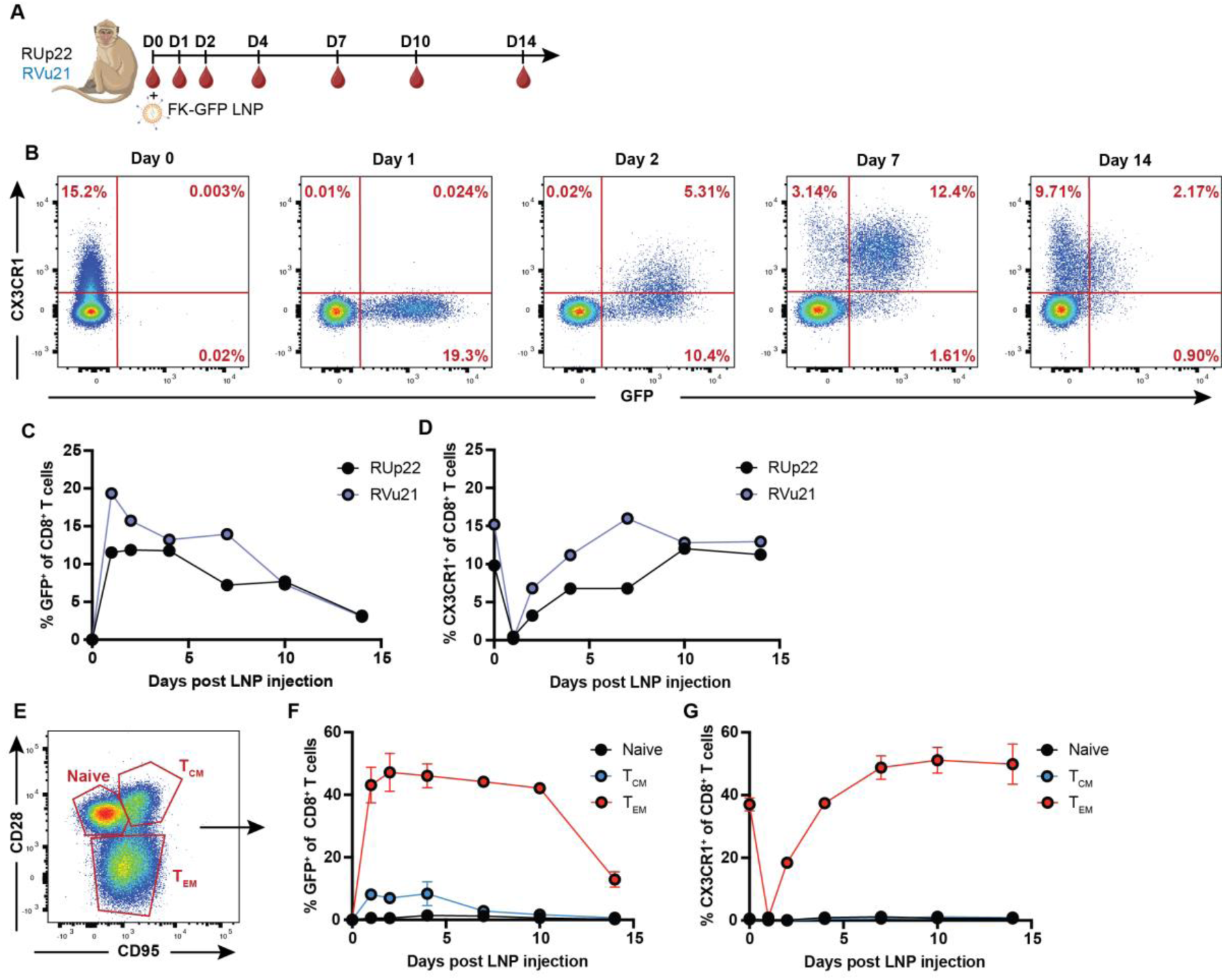
Fractalkine-conjugated mRNA-LNP target rhesus macaque CX3CR1^+^ T_eff_ cells in vivo: (A) Experimental design for in vivo testing of fractalkine-conjugated GFP-encoding mRNA-LNP in rhesus macaques (0.7mg/kg dosage, 1:2 μg mRNA/μg fractalkine ratio, IV delivery). RUp22 and RVu21 are the animal identification names. (B) GFP expression in peripheral blood CX3CR1^+^ T_eff_ cells either at baseline (Day 0) or after LNP injection (Days 1-14) with results from one animal shown. (C) Percentage of peripheral blood CD8^+^ T cells expressing GFP or (D) CX3CR1 across all timepoints analyzed. (E) Memory subset identification of CD8^+^ T cells based on CD28/CD95 to define naïve (CD28^+^/CD95^-^), central memory (T_CM_; CD28^+^/CD95^+^), and effector memory (T_EM_; CD28^-^/CD95^+^). (F) Percentage of GFP^+^ peripheral blood CD8^+^ T cells from each memory subset at each timepoint. (G) CX3CR1 expression levels at each timepoint.

## In vivo manipulation of rhesus macaque CX3CR1^+^ T_eff_ cells

Intravascular CX3CR1^+^ T_eff_ cells lack CD62L expression and are therefore generally unable to enter or exert cytotoxicity in lymph nodes at steady state (*32*). As such, enabling their trafficking into secondary lymphoid tissues could have substantial therapeutic applications for cancer and tissue residing viral infections. As CD62L is normally cleaved upon activation and lymph node homing (*33, 34*), we generated an mRNA encoding a shedding-deficient human CD62L (hCD62Lmut) (*35*). Expression of hCD62Lmut was readily detectable on RM CX3CR1^+^ T_eff_ cells and CX3CR1^+^ NK cells after delivery of fractalkine-conjugated hCD62Lmut mRNA-LNP in vitro using a human-specific CD62L monoclonal antibody (fig S12A-C). We subsequently dosed two RMs with a single dose of fractalkine-conjugated hCD62Lmut-encoding mRNA-LNP and monitored hCD62Lmut expression in CX3CR1^+^ T_eff_ cells across four days in comparison to two untreated control NHPs (Fig 5A). Both treated animals showed hCD62Lmut expression on peripheral blood CD8^+^ T cells 24 hours post LNP treatment corresponding to pre-treatment CX3CR1 expression levels (Fig 5B-C and fig S13A). As expected, hCD62Lmut expression was largely restricted to the T_EM_ subset (Fig 5D and fig S13B), demonstrating effector-specific modification. We were further able to identify hCD62Lmut expression on CD8^+^ T cells from lymph node fine needle aspirates 24 hours post LNP injection (Fig 5E and fig S13C). These hCD62Lmut^+^ CD8^+^ T cells had high expression of granzyme B, and were distinct from the more naïve rhesus CD62L^+^ CD8^+^ T cells also present within the lymph node (Fig 5F-G). Similar results were also found for peripheral blood and lymph node NK cells (fig S14A-F). Collectively, these data demonstrate the ability of fractalkine-conjugated LNP to transiently reprogram CX3CR1^+^ T_eff_ cells to express the homing molecule CD62L in vivo. The presence of hCD62Lmut^+^ CD8^+^ T cells within the lymph node is promising as it may suggest the expression of hCD62Lmut enabled lymph node trafficking of peripheral blood CX3CR1^+^ T_eff_ cells. However, as the untreated NHP also had notable granzyme B^+^ CX3CR1^+^ CD8^+^ T cell populations within the lymph node (fig S13C), further experiments will be necessary to determine whether the hCD62Lmut^+^ granzyme B^+^ CD8^+^ T cells found in lymph node were the result of altered lymphocyte trafficking.

**Fig. 5.**
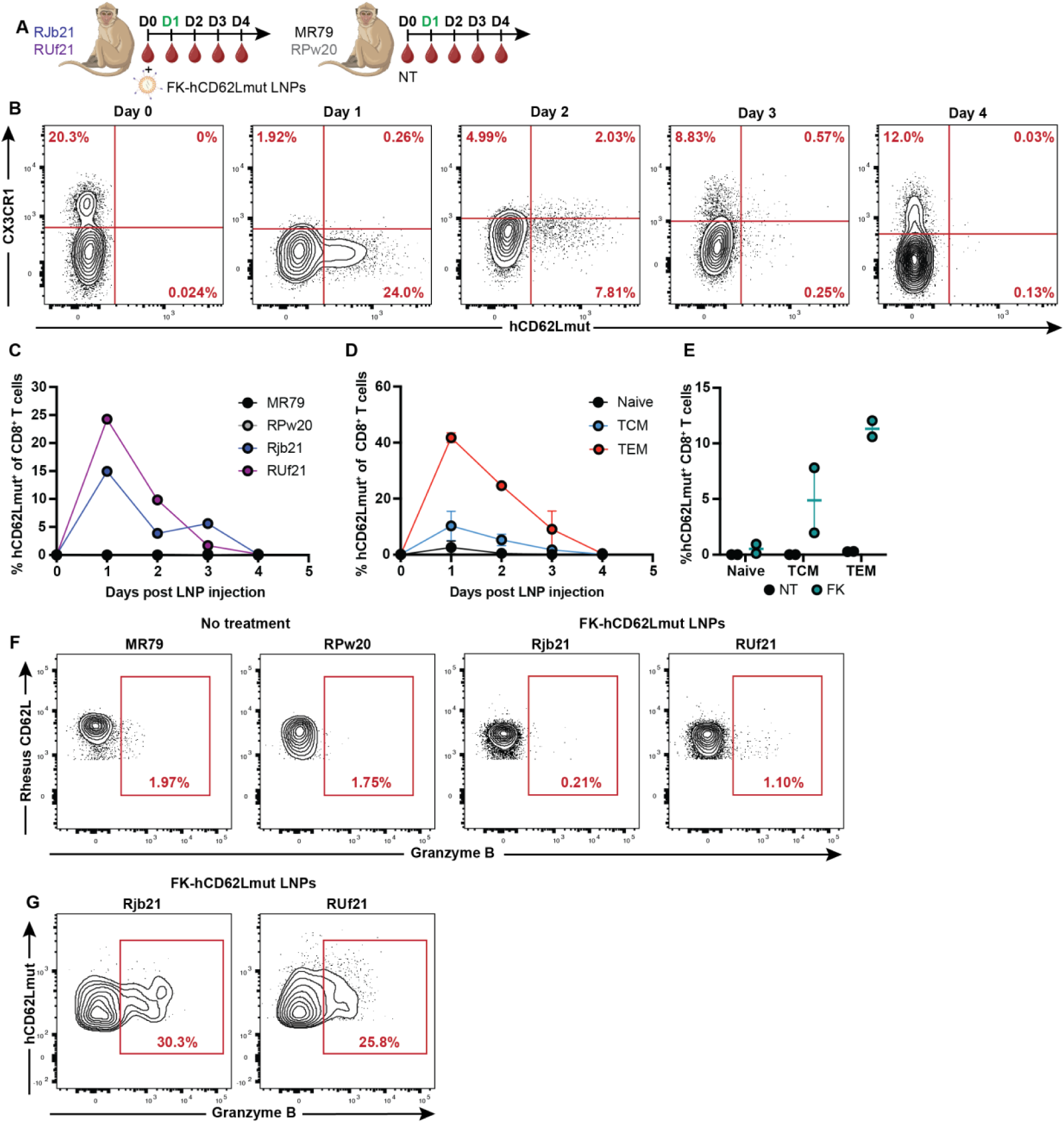
Fractalkine-conjugated mRNA-LNP drive hCD62Lmut expression in CX3CR1^+^ T_eff_ cells in vivo: (A) Experimental design for in vivo testing of fractalkine-conjugated human CD62L-encoding mRNA-LNP in rhesus macaques (0.5mg/kg dosage, 1:2 μg mRNA/μg fractalkine ratio). Rjb21, RUf21, MR79, and RPw20 are the animal identification names. NT = no treatment, FK = fractalkine. Green lettering represents fine need aspiration from lymph node. (B) hCD62Lmut expression in peripheral blood CX3CR1^+^ T_eff_ cells at baseline (Day 0) or after LNP injection (Days 1-4) with results from one animal shown. (C) Percentage of peripheral blood CD8^+^ T cells expressing hCD62Lmut across all timepoints analyzed. (D) Percentage of peripheral blood CD8^+^ T cells expressing hCD62Lmut in each CD8^+^ T cell memory subset across all timepoints analyzed. (E) Percentage of CD8^+^ T cells from lymph node expressing hCD62Lmut in each CD8^+^ T cell memory subset 24 hours post LNP injection. (F) Granzyme B expression in rhesus CD62L^+^ or (G) hCD62Lmut^+^ CD8^+^ T cells from lymph node 24 hours post LNP injection.

Overall, these experiments demonstrate that fractalkine, and more broadly, natural ligands, can be conjugated to mRNA-LNP as a novel strategy to target and reprogram distinct lymphocyte populations with high specificity. Our approach to readily and selectively in vivo reprogram only the CD8^+^ T cells with high cytotoxic potential provides a novel platform for mRNA-based therapies with a substantial list of potential therapeutic applications. The successful in vivo production and expression of hCD62Lmut on NHP CX3CR1^+^ T_eff_ cells represents a critical first step in expanding cytotoxic lymphocyte trafficking capabilities and highlights the ability of our system to transiently modify CX3CR1^+^ T_eff_ cells. Our platform is further enhanced by the capacity for sequential dosing of CX3CR1^+^ T_eff_ cells longitudinally with fractalkine-conjugated mRNA-LNP, which is highly favorable considering the transient nature of delivered mRNA expression. Furthermore, the consistency seen across targeting of human, mouse, and nonhuman primate cytotoxic effector CD8^+^ T cells by fractalkine-conjugated mRNA-LNP both in vitro and in vivo highlights the high translational potential of this targeting strategy and demonstrates in vitro targeting outcomes as an indicator for in vivo success. While future studies will be necessary to explore the efficacy of CD62L mRNA in driving lymphocyte trafficking, our strategy of conjugating fractalkine to mRNA-LNP holds tremendous promise as a targeted and scalable strategy to transiently reprogram CX3CR1^+^ T_eff_ cells for novel and/or enhanced biological functions.

## Acknowledgements

We would like to thank Garima Dwivedi and Mohamad-Gabriel Alameh for designing plasmids and preparing mRNA for LNP encapsulation, Christopher A. Hunter for providing the IL-2 mRNA-LNP, and Stephanie Ehnert, Sylvia West, and Stacie Weissman at the Emory National Primate Research Center for their assistance with all NHP studies. The data for this manuscript were generated in the Penn Cytomics and Cell Sorting Shared Resource Laboratory at the University of Pennsylvania and is partially supported by the Abramson Cancer Center NCI Grant (P30 016520). The research identifier number is RRID:SCR_022376. Lastly, we also thank Emily Cento, Zhilin Chen, Max A. Eldabbas, and Emileigh Maddox of the Human Immunology Core and the Division of Transfusion Medicine and Therapeutic Pathology at the Perelman School of Medicine at the University of Pennsylvania for providing de-identified human peripheral blood cells purified from healthy donor apheresis using StemCell RosetteSep™ kits. The HIC is supported in part by NIH P30 AI045008 and P30 CA016520. HIC RRID: SCR_022380.

## Funding

NIAID-funded ERASE HIV consortium UM1 AI-164562 (MP)

NIH Office of Research Infrastructure Programs P51OD11132 and the U42OD011023 (Emory National Primate Research Center)

NIH grants AI155577, AI115712, AI117950, AI108545, AI082630, AI149680, HL145754 (EJW)

Parker Institute for Cancer Immunotherapy funds (EJW)

The Mark Foundation (EJW)

NIAID-funded CIAVCR UM1 AI-169633 (MRB, DW, EFK)

BEAT-HIV consortium UM1 AI-164570 (MRB, DW, EFK)

Penn Measey Scholar in Molecular Medicine fund (EFK)

## Author contributions

MRB and VHW conceived of idea to conjugate fractalkine to mRNA LNP for T_eff_ targeting and express CD62L on cytotoxic T cells via mRNA-LNPs. ARC and EFK developed fractalkine-conjugation methods. ARC and AA performed fractalkine conjugation of LNP. ARC, MBP, EFK, VW, and MRB designed in vitro targeting experiments. ARC and MBP performed in vitro targeting experiments. ARC and SFN designed and performed all mouse experiments. JMLN, SDC and JH assisted with mouse IL-2 experiments. JMLN and VHW designed, performed, analyzed, and graphed single cell RNA sequencing experiments. EFK and DW designed mRNA constructs. ARC, AA, and JS produced mRNA. HN performed quality control experiments on mRNA. RR encapsulated all mRNA into LNP. ARC, MRB, EFK, MP, MS, and JH designed in vivo NHP experiments. MS performed flow cytometry for all NHP experiments. RLS and JW provided veterinary management of the NHP studies performed at Emory. MRB supervised all in vitro experiments. MRB and EJW supervised all mouse experiments. MRB and MP supervised all NHP experiments. MRB, EJW, DW, EFK, and MP provided funding. ARC prepared all figures and wrote manuscript.

## Data and materials availability

All data are available in the main manuscript or the supplementary materials. Any requests for materials should be addressed to ARC or MRB.

## Supplementary Materials

## Materials and Methods

### Mice

Female C57BL/6 mice between the ages of 6-8 weeks were purchased from Charles River (NCI) and maintained in SPF facility at the University of Pennsylvania. All conducted experiments were carried out in accordance with approval from the University of Pennsylvania UPenn Institutional Animal Care and Use Committee (IACUC).

### Human cell samples

Lymphocytes were harvested from 20 deidentified healthy donors via apheresis by the Human Immunology Core at the Perelman School of Medicine at the University of Pennsylvania. PBMCs were purified into CD8^+^ T cells and total T cell suspensions and provided by the Core using StemCell RosetteSep™ kits.

### Nonhuman primates

All animal experiments were conducted following guidelines established by the Animal Welfare Act and by the NIH’s Guide for the Care and Use of Laboratory Animals, Eighth edition. All the procedures were approved by the Emory University Institutional Animal Care and Use Committee (IACUC – Protocol number 202400000033). Animal care facilities at Emory National Primate Research Center are accredited by the U.S. Department of Agriculture (USDA) and the Association for Assessment and Accreditation of Laboratory Animal Care (AAALAC) International.

### mRNA synthesis and LNP encapsulation

Codon optimized inserts encoding eGFP, IL-2, and human CD62Lthat incorporates the TACE resistant CD62P transmembrane domain (hCD62Lmut) were cloned into an IVT expression vector as previously described (*44*). Low-endotoxin maxi preps were provided by Genscript (Piscataway, NJ). Nucleoside modified mRNA was produced from linearized DNA templates (restriction endonucleases from NE Biolabs) using the Megascript T7 Transcription Kit (Life Technologies) as previously described (*45*). Uridine was completely substituted for N1-methylpseudouridine (Trilink) and co-transcriptional capping was performed using AG 3OMe CleanCap (Trilink). Following synthesis, mRNA was purified of double stranded RNA products using cellulose purification (*46*). All mRNAs were assessed for degradation and appropriate size using gel electrophoresis and stored at -20C. mRNA was encapsulated into 4-component LNP using a self-assembly process as previously described (*19*). Briefly, an ethanolic mixture of component lipids (an ionizable cationic lipid, distearoylphosphatidylcholine, cholesterol and a polyethylene glycol (PEG) lipid) was mixed with the aqueous mRNA solution at an acidic pH. The ionizable cationic and PEG lipids and LNP composition are described in patent application WO 2017/004143A1. mRNA-LNP were assessed by dynamic light scattering using a Malvern Zetasizer and a Ribogreen assay for size, PDI, and encapsulation efficiency. All particles were ∼70nm in size and encapsulation efficiency was ∼95%.

### Fractalkine-conjugation of mRNA-LNP

Recombinant full length soluble human fractalkine containing both the chemokine domain and mucin stalk [Gln25-Arg339 (Ser199Asn)] (Genscript) or a truncated mouse fractalkine containing only the chemokine domain (Prospec Bio) was conjugated to mRNA-LNP using adapted methods from previous studies (*18, 19, 26, 47, 48*). Briefly, the recombinant fractalkine was functionalized via SATA (N-succinimidyl S-acetylthioacetate) to introduce sulfhydryl groups on cysteine residues present within the protein sequence. Hydroxylamine (0.5M) was used to deprotect SATA following purification from unreacted components via Zeba desalting spin columns. To test for impact of fractalkine amount, simultaneously, micelles composed of mPEG-maleimide were introduced into mRNA-LNP using a post-insertion technique. The functionalized fractalkine was then incubated with the mRNA-LNP, enabling the fractalkine to conjugate to the maleimide. Following conjugation, the fractalkine-conjugated mRNA-LNP were purified using Sepharose CL-4B-packed filtration columns and concentrated using 10kDa concentrators (Cytiva). The concentration of the mRNA-containing fractalkine-conjugated LNP was determined by performing a Quant-iT RiboGreen RNA assay. Conjugated LNP were stored at 4°C until usage.

### In vitro testing of fractalkine-conjugated mRNA-LNP

Cell suspensions of fresh or frozen purified human primary CD8^+^ T cells only, a mixture of CD8^+^ T cells and CD4^+^ T cells (total T cells), or PBMCs from healthy donors were plated in either RPMI containing Penicillin, Streptomycin and L-glutamine with 10% FBS or with 15% FBS and 30U/ml human IL-2 at a concentration of 1×10^6^cells/mL. Fractalkine-conjugated mRNA-LNP at dosages ranging from 0µg-1µg mRNA-LNP (as measured by RNA content) per million cells were added to the cell cultures and gently mixed. The LNP cell culture mixtures were incubated for times ranging from 16-72 hours at 37°C prior to collection and staining for flow cytometry.

### Flow cytometry for human samples

For in vitro human lymphocyte analysis, chemokine receptor staining was performed at 37°C for 15 minutes, followed by immediate addition of Live/Dead Aqua staining for 10 minutes at room temperature. Antibody cocktail for surface markers was then added for 20 minutes at room temperature. For extracellular staining only, cells were then washed with FACS buffer (PBS supplemented with 1% FBS) and resuspended in PBS. For conditions involving intracellular staining, after washing the cells with FACS buffer, the cells were pre-fixed with a 2% paraformaldehyde (PFA)/PBS mixture for 2-5 minutes prior to intracellular staining using a BD Cytofix/Cytoperm kit for one hour at room temperature. Antibodies used include APC-Cy7 Anti-Human CCR7 clone G043H7 (Biolegend), BV650 Anti-Human CX3CR1 clone 2A9-1 (Biolegend), BUV805 Anti-Human CD3 clone UCHT1 (BD), BV510 Anti-Human CD19 clone HIB19 (Biolegend), BUV737 Anti-Human CD27 clone 0323 (BD), BB790 Anti-Human CD4 clone SK3 (BD), BUV496 Anti-Human CD8 clone RPA-T8 (BD), PE-Dazzle 594 Anti-Human CD57 clone HNK-1 (Biolegend), BUV563 Anti-Human CD45RA clone HI100 (BD), BV510 Anti-Human CD14 clone M5E2 (Biolegend), APC Anti-Human Perforin clone B-D48 (Biolegend), AF700 Anti-Human Granzyme B clone GB11 (BD), BV785 Anti-Human Ki67 clone B56 (BD), PE-Cy7 Anti-Human Perforin clone B-D48 (Biolegend), BV421 Anti-Human Perforin clone B-D48 (Biolegend), PE-CF594 Anti-Human Granzyme B clone GB11 (BD), AF700 Anti-Human TCR Va7.2 (Biolegend), BUV395 Anti-Human HLA-DR clone G46-6 (BD), BUV615 Anti-Human CD16 clone 3G8 (BD), BV480 Anti-Human CD14 clone MP9 (BD), BV570 Anti-Human CD56 clone HCD56 (Biolegend), BV605 Anti-Human CD69 clone FN50 (Biolegend), BV711 Anti-Human CD38 clone HIT2 (Biolegend), BV785 Anti-Human CD8 clone RPA-T8 (Biolegend), and PE-Cy5 Anti-Human CD161 clone DX12 (BD).

### Single-Cell 3’ RNA Sequencing Sample Processing and Library Generation

Between 667,000 and 1 million PBMCs from four healthy donors were plated in complete R10 medium (RPMI ^+^ 10% FBS ^+^ 1% penicillin/streptomycin ^+^ 2 mM L-glutamine) supplemented with 30U/mL human IL-2 at a concentration of 1 × 10⁶ cells/mL. Cells were treated with either 0.5μg per million cells of unconjugated mRNA-LNP GFP, fractalkine-conjugated mRNA-LNP GFP, or PBS, and gently mixed. The LNP-cell culture mixtures were incubated overnight (∼24 hours) at 37°C with 5% CO₂. All subsequent cell preparation and antibody staining steps, including centrifugation, were performed at 10°C or on ice. After overnight incubation, cells were collected, centrifuged at 500 × g for 5 minutes, and resuspended in 1mL of ice-cold staining buffer (PBS ^+^ 2% BSA ^+^ 0.01% Tween-20). Cells were then spun again at 400 × g for 5 minutes and resuspended in 22.5μL of staining buffer. Next, 2.5μL of TruStain was added to each sample and incubated on ice for 10 minutes. Following this incubation, each donor sample was stained with a unique hashing antibody (HTO, TotalSeq-A Hashing, BioLegend) following the manufacturer’s instructions. The antibodies were diluted in 25μL of staining buffer and added to the cells for a final staining volume of 50μL. After a 30-minute incubation on ice, cells were washed three times with ice-cold staining buffer, counted, and pooled in the staining buffer, targeting ∼150,000 cells per sample. The pooled samples were then filtered through a 40μm FlowMi strainer (Sigma) and counted again before being diluted to the 10X Genomics recommended concentration. Samples were processed using the Chromium X platform (10X Genomics). Single-cell sequencing library preparations for the RNA modality followed the 10X Genomics User Guide *Chromium Next GEM Single Cell 3’ Reagents Guide v3.1 (Dual Index)* (CG000315 Rev E), with the exception of spiking in HTO additive primers at step 2.2 to amplify HTO sequences. Specifically, 1μL of 0.1μM HTO additive primer (sequence: GTGACTGGAGTTCAGACGTGTGCTCTTCCGAT*C*T) was added to each individual sample preparation. HTO single-cell sequencing library preparation followed the antibody manufacturer’s protocol (BioLegend) and was conducted as described by the Satija Lab on cite-seq.com. Briefly, at step 2.3d of the 10X protocol, 60μL of supernatant was collected and mixed with 140μL of SPRIselect reagent (Beckman Coulter), washed twice with 80% ethanol, and eluted with 45μL of EB buffer (Qiagen). For the HTO modality, antibody-derived oligomers were amplified and indexed by mixing 45μL of eluate with 50μL of KAPA HiFi HotStart ReadyMix (KAPA Biosystems) and 5μL of forward/reverse indexing primers (BioLegend). Post-amplification, HTO libraries were purified with 120μL SPRIselect cleanup and eluted with 30μL of EB buffer. All final libraries were quantified using a Qubit dsDNA HS Assay Kit (Invitrogen) and analyzed using a High Sensitivity D1000 DNA tape (Agilent) on a TapeStation D4200 (Agilent).

### Single-Cell 3’ RNA and HTO Library Sequencing

Sequencing was performed on the NovaSeq 6000 platform (Illumina) with a target of at least 25,000 reads per cell for RNA libraries and 1,000 reads per cell for HTO libraries. Libraries were denatured and diluted according to the *Illumina NovaSeq 6000 Protocol A: Pool and Denature Libraries for Sequencing (Standard Loading, Document #1000000106351 v03)*.

### Single-cell RNA sequencing analysis

Raw sequencing files were demultiplexed using cellranger mkfastq (10X Genomics) and bcl2fastq2 (Illumina). HTO files were then processed using the KITE pipeline with kallisto and bustools (*50*). Briefly, fastq files were aligned to a reference index with the TotalSeqA (BioLegend) hashtag barcodes used in the experiment. Resultant alignments were collated with corrected cell barcodes and UMIs to produce a cell by feature matrix. GEX/RNA files were processed using cellranger count (10X Genomics). HTO and GEX outputs were then analyzed in R (v4.1.1) using Seurat (*51*). Cells were quality control filtered (number of features > 200, number of features < 6000, and percent of counts pertaining to mitochondrial genes < 12.5%), dehashed (using MULTIseqDemux from the Seurat package), and filtered for doublets via the hashing antibodies. The dataset was then annotated using reference-based dataset on PBMCs (*52*). Differential gene expression CD8 Tem and NK cells was calculated using the function wilcoxaux from the presto package (https://github.com/immunogenomics/presto). Gene set enrichment analysis (GSEA) was then performed using the msigdbr and fgsea packages (*53, 54*). Graphs were made using ggplot2 and patchwork. All raw sequencing data have been deposited to the Gene Expression Omnibus (GEO) with the accession record GSE296681.

### Mouse LCMV Armstrong infection

Mouse LCMV Armstrong was propagated and assessed for viral titer as previously described (*36, 37*). Mice were then infected intraperitoneally with 2×10^5^ plaque forming units of LCMV Armstrong.

### Tissue processing of mouse samples

For PBMC isolation, blood was first collected in a 4% sodium citrate solution. PBMCs were then isolated using Histopaque-1083-based gradient centrifugation and washed in RPMI containing 1% FBS prior to cell staining. For serum collection, blood was collected in 1.5mL microcentrifuge tubes and placed on ice. Samples were then spun down at 15,000rpm for 10 minutes at 4°C. Serum was then collected and stored at -80°C until usage in ELISA. For splenocyte collection, spleens were collected and placed in RPMI containing 1% FBS on ice. The spleens were then smashed, filtered through a 70µm cell strainer, and resulting splenocytes were washed with RPMI containing 1% FBS prior to cell staining. Red blood cells were lysed with ACK Lysis Buffer.

### Flow cytometry for mouse samples

For mouse cell analysis, PBMCs and splenocytes were stained with Zombie Yellow Fixable Viability Dye for 30 minutes at 4°C prior to staining with antibody cocktail for extracellular surface markers for 30 minutes at 4°C. Both human and mouse samples were acquired on a Symphony A5 and data were analyzed using FlowJo software. Antibodies used include BD OptiBuild BV750 Hamster Anti-Mouse CD27 (BD), BUV737 Rat Anti-Mouse CD127 Clone SB/199 (BD), BUV805 Rat Anti-Mouse CD8a Clone 53-6.7 (RUO) (BD), BV605 Anti-Mouse CX3CR1 clone SA011F11 (BioLegend), BD OptiBuild BV421 Rat Anti-Mouse CD94 (BD), BD OptiBuild BUV615 Hamster Anti-Mouse KLRG1 (BD), BUV395 Rat Anti-Mouse CD44 clone IM7 (BD), BUV563 Rat Anti-Mouse CD4 clone GK1.5 (BD), GP33 tetramer-PE

### Mouse in vivo testing of fractalkine-conjugated mRNA-LNP

For studies involving mRNA-LNP encoding fluorescent proteins, C57BL/6 mice were infected with LCMV Armstrong. Between days 8-33 post infection the mice were then retro-orbitally injected with either PBS, unconjugated GFP, unconjugated tdTomato, fractalkine-conjugated GFP or fractakine-conjugated tdTomato LNP. For all studies aside from dose escalation study, LNP were given at a dosage of 10µg in 200µL PBS solution. The dose was calculated based on the mRNA content. PBS-treated mice were given 200µL PBS. Mice were randomly distributed amongst cages prior to any treatment condition. The µg mRNA/µg fractalkine ratios of the mouse fractalkine-conjugated mRNA-LNP ranged from 1:3 to 1:0.75 depending on the study. For testing mouse fractalkine-conjugated IL-2-encoding mRNA-LNP, C57BL/6 mice were infected with LCMV Armstrong. Eleven days post infection the mice were randomly distributed amongst cages and given either 200µL PBS, or 6.5µg in 200µL PBS unconjugated IL-2 LNP or 6.5µg in 200µL PBS fractalkine-conjugated IL-2 LNP.

### Flow cytometry analysis of IL-2 secretion in splenocytes

Three hours after PBS, unconjugated or mouse fractalkine-conjugated IL-2 LNP injection, the mice were sacrificed and splenocytes were isolated. An IL-2 secretion assay (Miltenyi Biotec) was performed (*49*). Briefly, splenocytes were washed twice and resuspended in cold buffer (PBS supplemented with 0.5% bovine serum albumin (BSA) and 2 mM EDTA). Cytokine Catch Reagent was then added to the cells and incubated on ice for 5 minutes. The cells were then incubated in a closed tube containing warmed RPMI media supplemented with 10% FBS under continuous rotation for two hours at 37°C. The cells were then washed twice and resuspended in cold buffer. Cytokine Detection Antibody was then added to the cells, along with an antibody cocktail for extracellular staining, and incubated for 10 minutes on ice. The cells were then washed twice and acquired on flow cytometer.

### Flow sorting and culture of splenic CD8^+^ T cells

Following flow staining, pooled spleen samples (i.e., mice treated with PBS, unconjugated LNP, or fractalkine-conjugated LNP) were sorted using the Aurora CS system, simultaneously collecting CX3CR1^+^ and CX3CR1^-^ CD8^+^ T cells. After sorting, cells were spun down and resuspended in T cell media (Minimum Essential Media containing 10% fetal bovine serum, 2 mM L-glutamine, penicillin/streptomycin, 12.5 mM HEPES, and β-mercaptoethanol). Approximately 2.5e^5^ cells from each condition were plated into one well of a 24-well plate in a final volume of 550µl of T cell media and left at 37°C in 5% CO_2_ for three hours. Following the incubation, the media from each well was collected and any residual cells were spun down. The supernatant was split into two 250µl aliquots and frozen at -80°C until analyzing via ELISA.

### IL-2 ELISA

ELISA to test for presence of IL-2 in mouse serum and sorted splenic CD8^+^ T cell culture supernatants was carried out using an ELISA MAX™ Standard Set (Biolegend). Briefly, Nunc™ MaxiSorp™ ELISA Plates were coated with Capture Antibody overnight at 4°C. The following day, the plates were washed with three times with PBS-T (0.10% tween/PBS) and subsequently blocked using a 10% FBS/PBS buffer solution. The plates were then washed, and serially diluted mouse samples and control antibodies were added and incubated for two hours while shaking at 500rpm. The plates were then washed, and detection antibody was added for one hour with shaking. The plates were then washed and an avidin-HRP solution was added and incubated for 30 minutes with shaking. The plates were washed and were developed using a TMB Substrate Set (Biolegend) for 20 minutes. The reaction was stopped by adding in 2N H_2_SO_4_ and the plates were read for absorbance at 450nm.

### Nonhuman Primate in vivo testing of fractalkine-conjugated mRNA-LNP

The fractalkine-conjugated GFP LNP experiment included two specific-pathogen free Indian Rhesus macaques (RM; *Macaca mulatta*), single-housed in an animal BSL-2 facility at Emory National Primate Research Center, Atlanta, GA [1 female (RUp22), 1 male (RVu21), *Mamu*-A*01-, *Mamu*-B*08^−^, *Mamu*-B*17^−^, 2-3 years at the start of the study]. Both animals received a 0.7 mg/kg dose of fractalkine-conjugated GFP mRNA-LNP. The fractalkine-conjugated hCD62Lmut experiment included four specific-pathogen free Indian Rhesus macaques (RM; *Macaca mulatta*), single-housed in an animal BSL-2 facility at Emory National Primate Research Center, Atlanta, GA [4 males (MR79, RJb21, RPw20, RUf21) *Mamu*-A*01-, *Mamu*-B*08^−^, *Mamu*-B*17^−^, 5-6 years at the time of first LNP infusion]. RM were infected intravenously with 10,000 IU of barcoded SIVmac239M and beginning at day 14 post infection received a daily subcutaneous antiretroviral therapy regimen of FTC (40 mg/kg), TDF (5.1 mg/kg) and DTG (2.5 mg/kg), that was maintained for 57 weeks before ATI. Two RM (RJb21, RUf21) received one intravenous 0.5mg/kg dose of fractalkine-conjugated hCD62Lmut mRNA-LNP four days post antiretroviral therapy interruption.

### Tissue processing of nonhuman primate samples

PBMCs and lymph node fine needle aspirates (FNAs) were collected longitudinally and processed as previously described (*38–41*). Briefly, PBMCs were isolated from whole blood by density gradient centrifugation with 90% ficoll. FNAs were harvested from one node using an ultrasound guided needle, and lymph node mononuclear cells were isolated via centrifugation (MR79 right axillary LN, RJb21 right inguinal LN, RPw20 right axillary LN, RUf21 right axillary LN). Mononuclear cells were counted for viability using a Countess II Automated Cell Counter (Thermo Fisher) with trypan blue stain. All samples were processed, stained and analyzed by flow cytometry immediately after collection.

### Flow cytometry for nonhuman primate samples

Multiparametric flow cytometry was performed using standard procedures on fresh PBMCs and mononuclear cells derived from LN aspirates using anti-human monoclonal antubidies previously shown to be cross-reactive in RM (*39, 42, 43*). The following antibodies were utilized for the staining panel to assess GFP expression: anti-CX3CR1-PE (clone K0124E1, 0.5μL – Biolegend cat.#355704), anti-CCR7-BB700 (clone 3D12, 5μL – BD cat.#566437), anti-CD16-BV421 (clone 3G8, 5μL – BD cat.#562874), anti-CD95-BV605 (clone DX2, 5μL – Biolegend cat.#305628), anti-HLA-DR-BV650 (clone L243, 5μL – Biolegend cat.#307650), anti-CD20-BUV786 (clone 2H7, 5μL – BD cat.#568713), anti-NKG2A-PE-Cy7 (clone Z199, 0.5μL – Beckman Coulter cat.#B10246), anti-CD3-BUV395 (clone SP34-2, 2.5μL – BD cat.#564117), anti-CD8-BUV496 (clone RPA-T8, 2.5μL – BD cat.#612942), anti-CD28-BUV737 (clone CD28.2, 2.5μL – BD cat.#612815), anti-CD14-BUV805 (clone M5E2, 5μL – BD cat.#612902), anti-CD4-APC-Cy7 (cone OKT4, 2.5μL – Biolegend cat.#317418), BD Horizon Brilliant Stain Buffer Plus (10μL – BD cat.#566385), Fixable Viability Stain 700 (0.5μL – BD cat.#564997). Acquisition was performed on a minimum of 100,000 live cells on a A5 Symphony flow cytometer (BD Biosciences) driven by BD FACSDiva software. Acquired data was analyzed using FlowJo v.10.10. The following antibodies were utilized for the staining panel to assess CD62L expression: anti-CX3CR1-PE (clone K0124E1, 0.5μL – Biolegend cat.#355704), anti-CCR7-BB700 (clone 3D12, 5μL – BD cat.#566437), anti-CCR5-BUV661 (clone 3A9, 5μL – BD cat.#750299), anti-CXCR5-PE-Cy5 (clone MU5UBEE, 5μL – Invitrogen cat.#15-9185-42), anti-CD62L-AlexaFluor488 (clone DREG-56, 0.15μL – Biolegend cat.#304816), anti-CD62L-BV750 (clone SK11, 5μL – BD OptiBuild cat.#747199), anti-CD16-BV421 (clone 3G8, 5μL – BD cat.#562874), anti-CD95-BV605 (clone DX2, 5μL – Biolegend cat.#305628), anti-HLA-DR-BV650 (clone L243, 5μL – Biolegend cat.#307650), anti-CD20-BUV786 (clone 2H7, 5μL – BD cat.#568713), anti-NKG2A-PE-Cy7 (clone Z199, 0.5μL – Beckman Coulter cat.#B10246), anti-CD3-BUV395 (clone SP34-2, 2.5μL – BD cat.#564117), anti-CD8-BUV496 (clone RPA-T8, 2.5μL – BD cat.#612942), anti-CD28-BUV737 (clone CD28.2, 2.5μL – BD cat.#612815), anti-CD14-BUV805 (clone M5E2, 5μL – BD cat.#612902), anti-CD4-APC-Cy7 (cone OKT4, 2.5μL – Biolegend cat.#317418), anti-CD69-BV711 (clone FN50, 5μL – BD cat.#563836), anti-PD-1-PE/Dazzle594 (clone EH12.2H7, 5μL – Biolegend cat.#329940), anti-CD45-BUV563 (clone D058-1283, 5μL – cat.#741414), BD Horizon Brilliant Stain Buffer Plus (10μL – BD cat.#566385), Fixable Viability Stain 700 (0.5μL – BD cat.#564997). Samples underwent fixation and permeabilization with Tonbo Foxp3/Transcription Factor Staining Buffer Kit (Cytek Bio, cat.#TNB-0607-KIT) for 45 min at 4 °C. Intracellular staining was performed for 30 minutes at room temperature with a cocktail containing anti-Granzyme B-APC (clone GB11, 2.5μL – Beckman Coulter cat.#GRB05), and anti-Ki-67-BV480 (clone B56, 5μL – BD cat.#566109). Acquisition was performed on a minimum of 100,000 live cells on a A5 Symphony flow cytometer (BD Biosciences) driven by BD FACSDiva software. Acquired data was analyzed using FlowJo v.10.10.

---

**Fig. S1.**
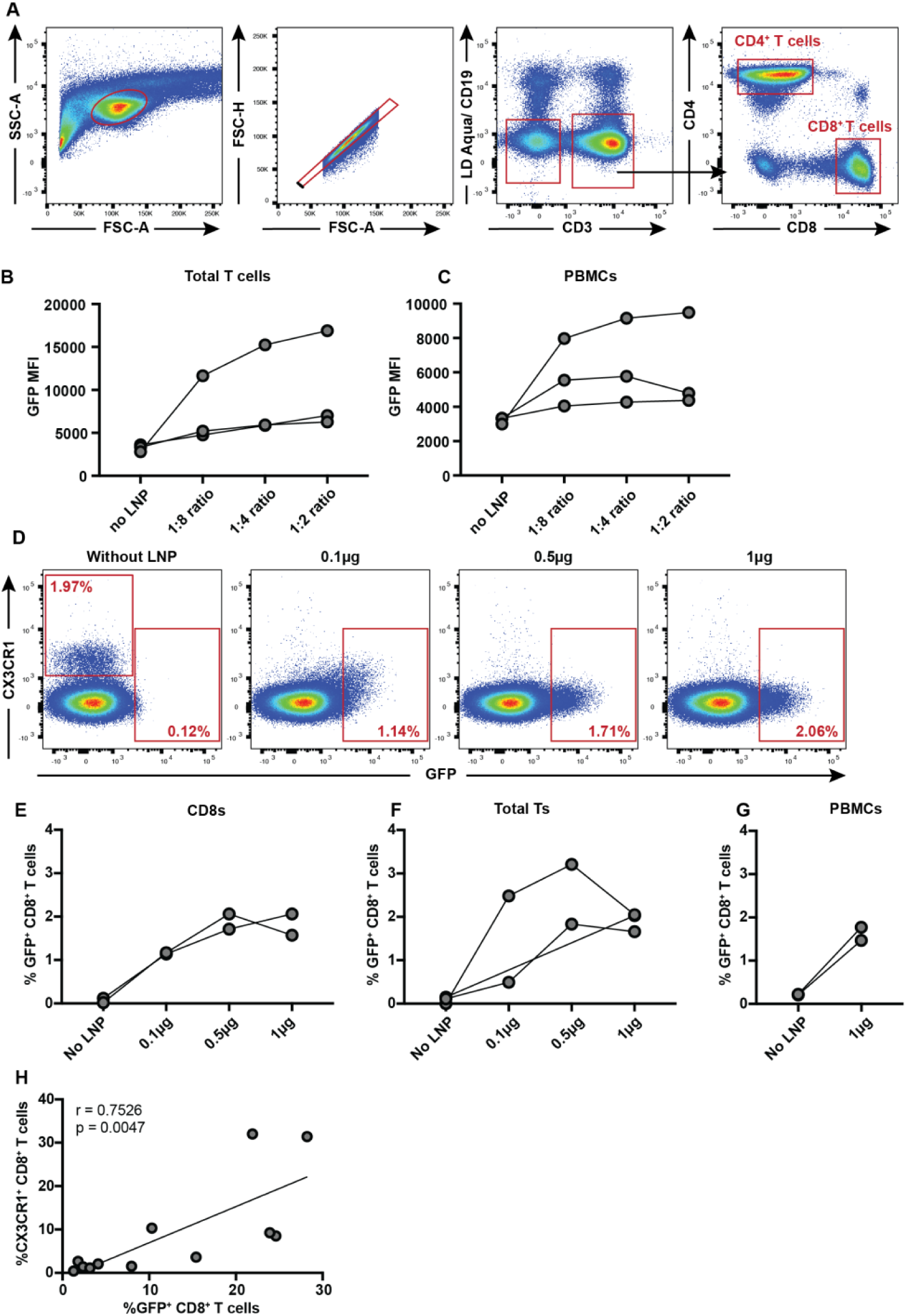
Fractalkine-conjugated LNP target CX3CR1^+^ T_eff_ cells in vitro: (A) Representative plots from a PBMC sample showing the gating strategy for CD8^+^ T cell and CD4^+^ T cell analysis. (B) Median fluorescence intensity of GFP in CD8^+^ T cells from total T cell and (C) PBMC conditions after a 65-hour incubation with or without human fractalkine-conjugated mRNA-LNP with mRNA-to-fractalkine μg ratios of 1:8, 1:4, and 1:2 at a dose of 0.5µg mRNA/million cells. Individual data points represent different donors. (D) Flow cytometry analysis of purified human primary CD8^+^ T cells incubated for 65 hours with or without fractalkine-conjugated GFP-encoding mRNA-LNP (1:8 ratio) at dosages ranging from 0.1-1µg mRNA/million cells. Data from one donor shown. (E) Percentage of CD8^+^ T cells expressing GFP from purified CD8^+^ T cells represented in (D), (F) total T cells and (G) PBMC. Individual data points represent different donors. (H) Correlation of the percentage of CX3CR1^+^ T_eff_ cells without LNP treatment and percentage of GFP^+^ CD8^+^ T cells after LNP treatment for all donors tested in fractalkine ratio experiment, dose-response experiment, and time-course experiment for fractalkine-conjugated LNP made at 1:2 mRNA/ fractalkine ratio. Pearson correlation coefficiency test used.

**Fig. S2.**
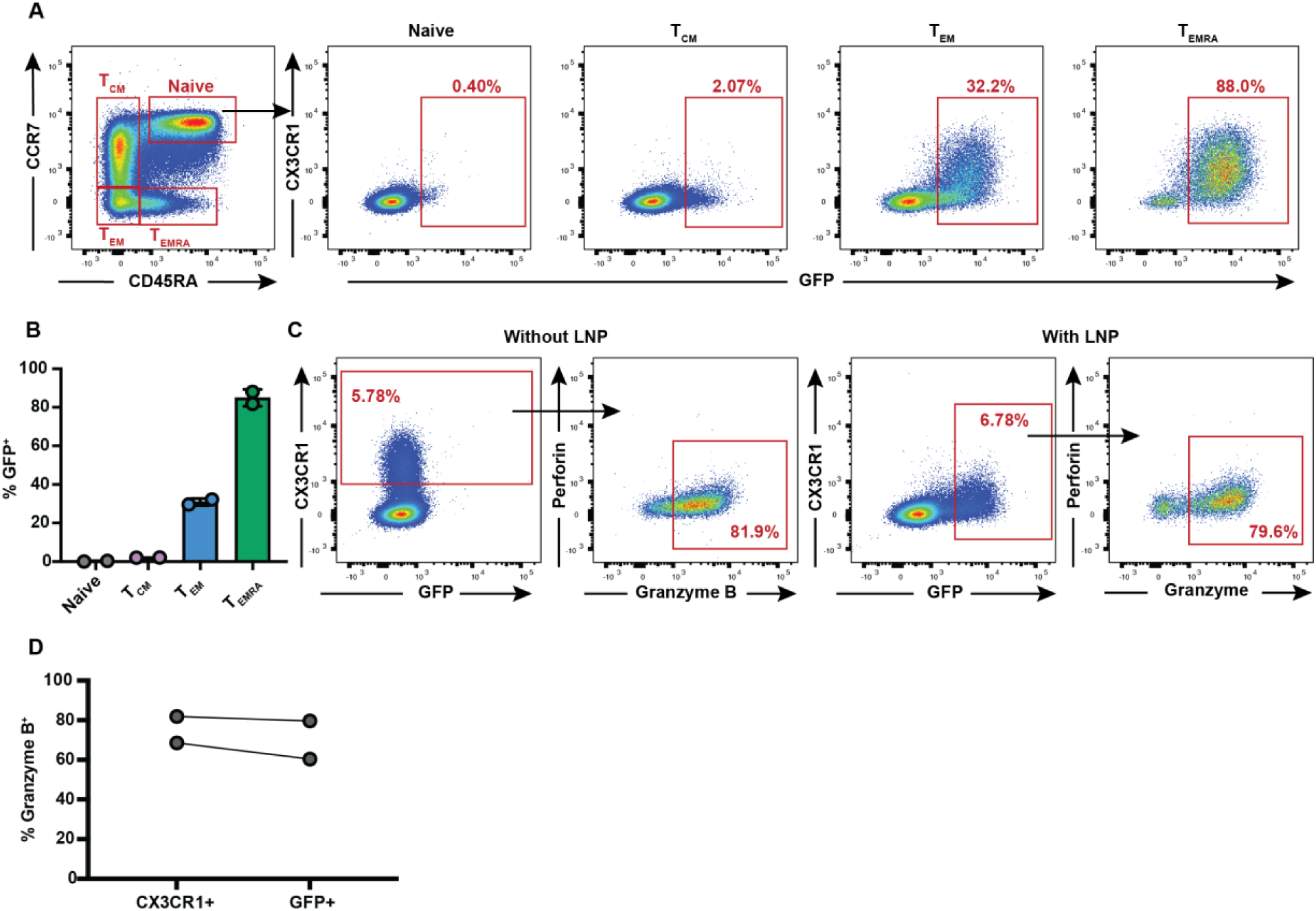
Fractalkine-conjugated LNP target CX3CR1^+^ CD4^+^ T cells in vitro: (A) Memory subset analysis of GFP expression patterns in canonical CD4^+^ T cell memory subsets, including naïve (CCR7^+^/CD45RA^+^), central memory (T_CM_; CCR7^+^/CD45RA^-^), effector memory (T_EM_; CCR7^-^/CD45RA^-^) and terminally differentiated effectors (T_EMRA_; CCR7^-^/CD45RA^+^), in total T cells after 65-hour incubation with fractalkine-conjugated GFP-encoding mRNA-LNP at 1:2 RNA/Fractalkine ratio. One representative donor shown. (B) GFP expression in each CD4^+^ T cell memory subset from (A) in donors with notable cytotoxic CD4^+^ T cell populations. (C) Perforin and granzyme B expression in CX3CR1^+^ CD4^+^ T cells from PBMC incubated for 65 hours alone (left two panels) or in GFP^+^ CD4^+^ T cells treated with fractalkine-conjugated GFP-encoding mRNA-LNP at a dose of 0.5µg mRNA/million cells (1:2 ratio) for 65 hours (right two panels). One representative donor shown. (D) Percentage of CX3CR1^+^ or GFP^+^ CD4^+^ T cells from PBMC expressing granzyme B as shown in (C), from all donors with notable cytotoxic CD4^+^ T cell populations.

**Fig. S3.**
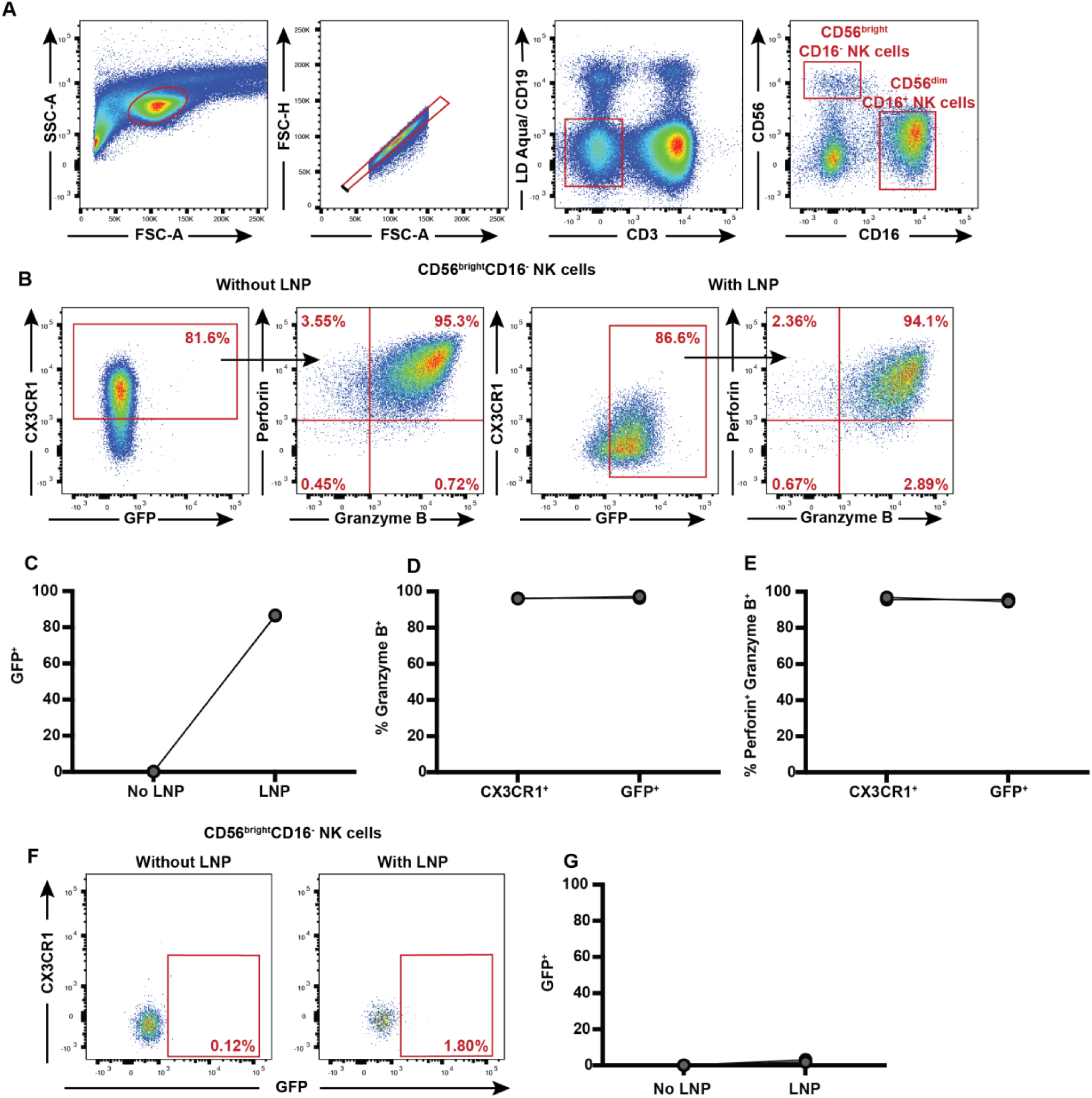
Fractalkine-conjugated LNP target CX3CR1^+^ NK cells in vitro: (A) Flow cytometry gating strategy for analyzing NK cells. (B) Expression of perforin and granzyme B in CX3CR1^+^ CD56^dim^CD16^+^ NK cells from PBMC incubated for 65 hours without LNP (left two panels) or in GFP^+^ CD56^dim^CD16^+^ NK cells from PBMC that were incubated with fractalkine-conjugated GFP-encoding mRNA-LNP at a dose of 0.5µg mRNA/million cells (1:2 ratio) for 65 hours (right two panels). One representative donor shown. (C) Percentage of CD56^dim^CD16^+^ NK cells expressing GFP as shown in (B) for all donors tested. (D) Percentage of CX3CR1^+^ or GFP^+^ CD56^dim^CD16^+^ NK cells from PBMC expressing granzyme B or (E) both perforin and granzyme B as shown in (B) for all donors tested. (F) Expression of CX3CR1 and GFP in CD56^bright^CD16^-^ NK cells from PBMC incubated for 65 hours without LNP (left panel) or with fractalkine-conjugated GFP-encoding mRNA-LNP at a dose of 0.5µg mRNA/million cells (1:2 ratio) for 65 hours (right panel) with results from one representative donor shown. (G) Percentage of GFP^+^ CD56^bright^CD16^-^ NK cells from (F) for all donors tested.

**Fig. S4.**
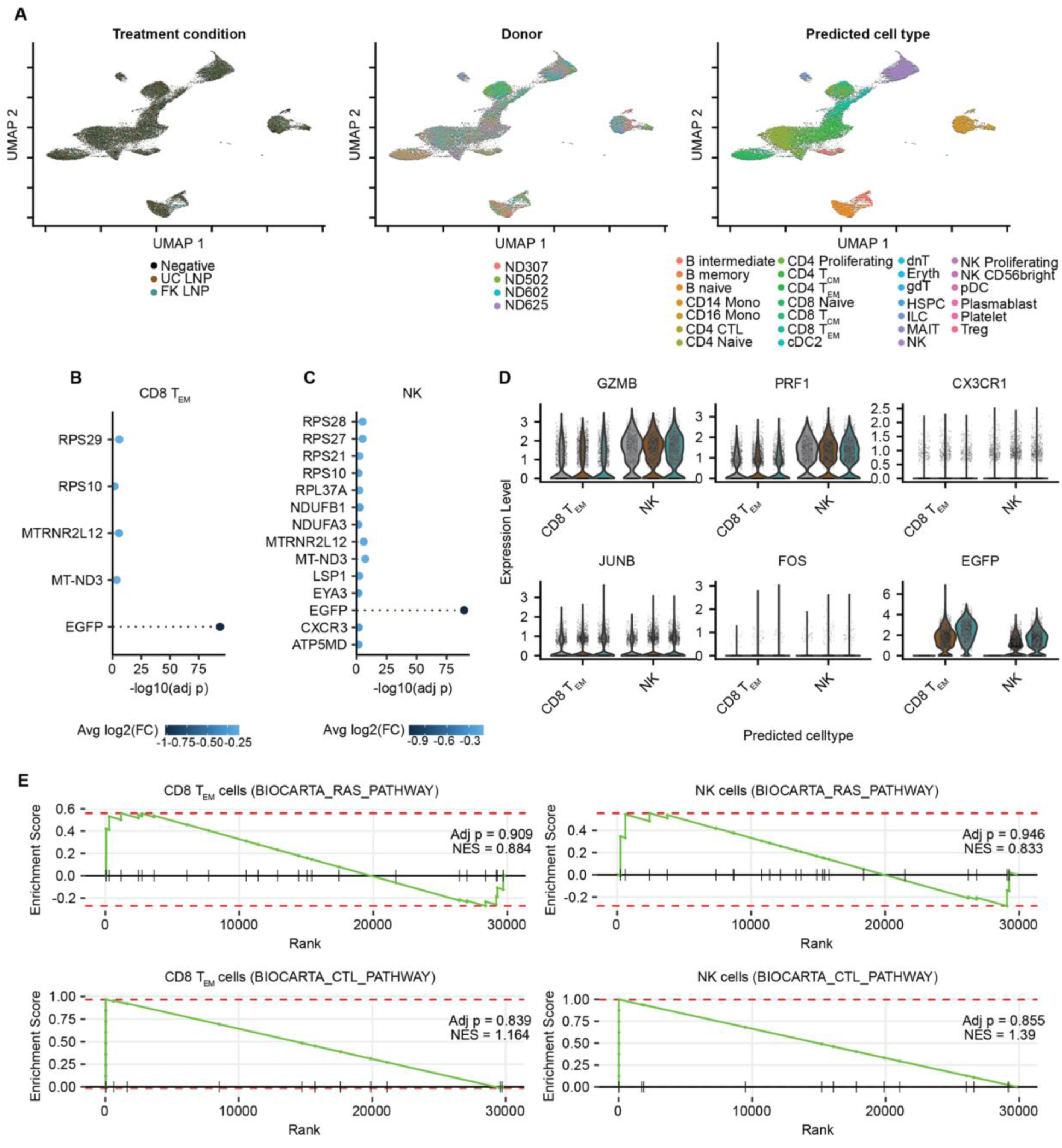
Single-Cell 3’ RNA Sequencing of fractalkine-conjugated LNP-treated CX3CR1^+^ cells: (A) UMAP representations of the single cell RNA sequencing dataset colored by treatment condition (left panel), donor (middle panel), and predicted cell type (right panel). (B) Differentially expressed genes in CD8^+^ TEM cells and (C) NK cells with negative values representing enriched genes in fractalkine-conjugated LNP treatment condition in comparison to unconjugated LNP condition across all donors combined. (D) Expression of granzyme B (top left panel), perforin-1 (top middle panel), CX3CR1 (top right panel), JunB (bottom left panel), FOS (bottom middle panel), and eGFP genes (bottom right panel) by CD8^+^ TEM cells and NK cells. Black = no treatment, brown = unconjugated LNP, teal = fractalkine-conjugated LNP. (E) Analysis of enrichment of either Ras signaling (top panels) or cytotoxic T lymphocyte (CTL) pathway signaling (bottom panels) in CD8^+^ TEM cells and NK cells after treatment with fractalkine-conjugated LNP in comparison to unconjugated LNP.

**Fig. S5.**
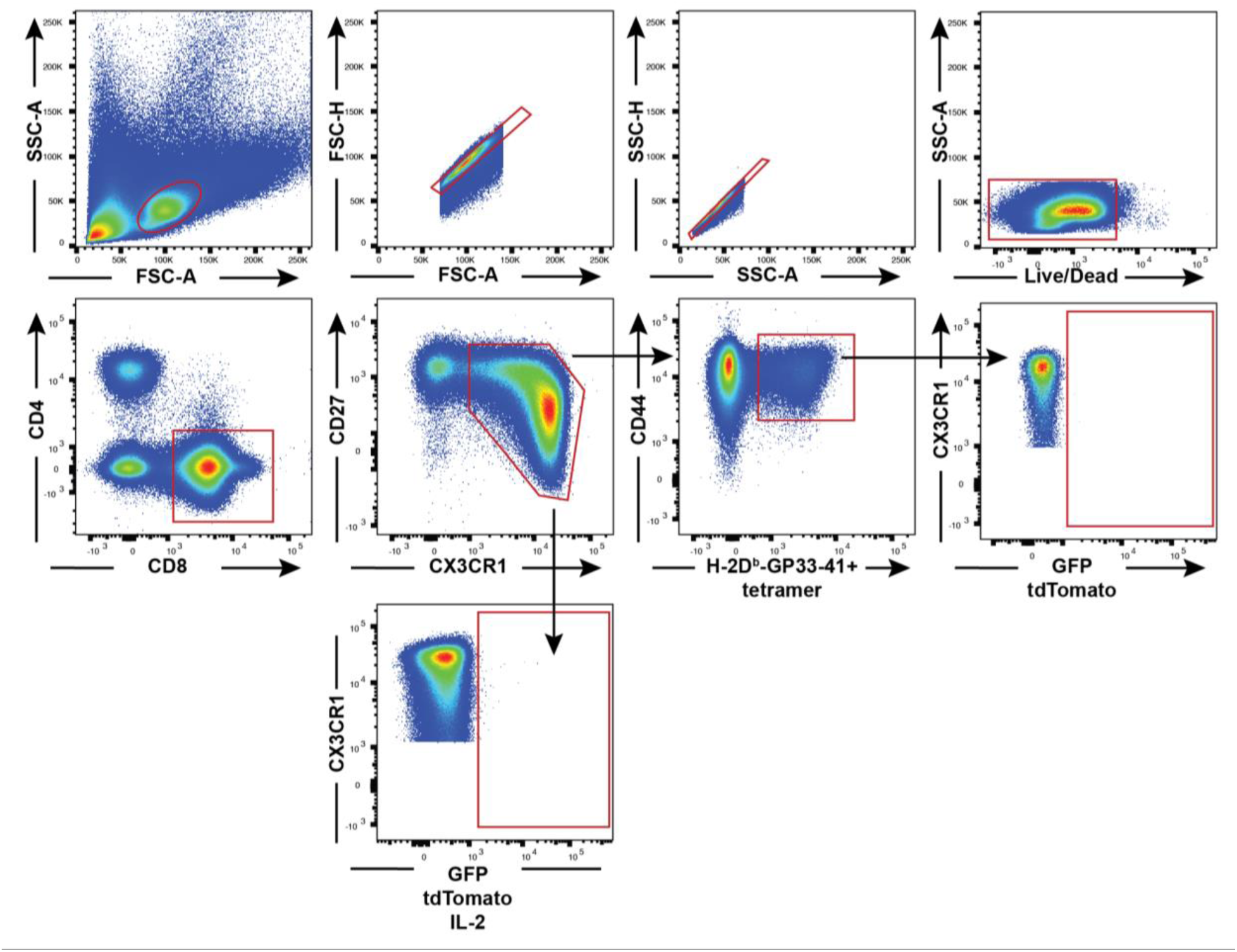
Flow cytometry panel used for analyzing mouse CD8^+^ T cells: Flow cytometry gating strategy for mouse CD8^+^ T cell analysis.

**Fig. S6.**
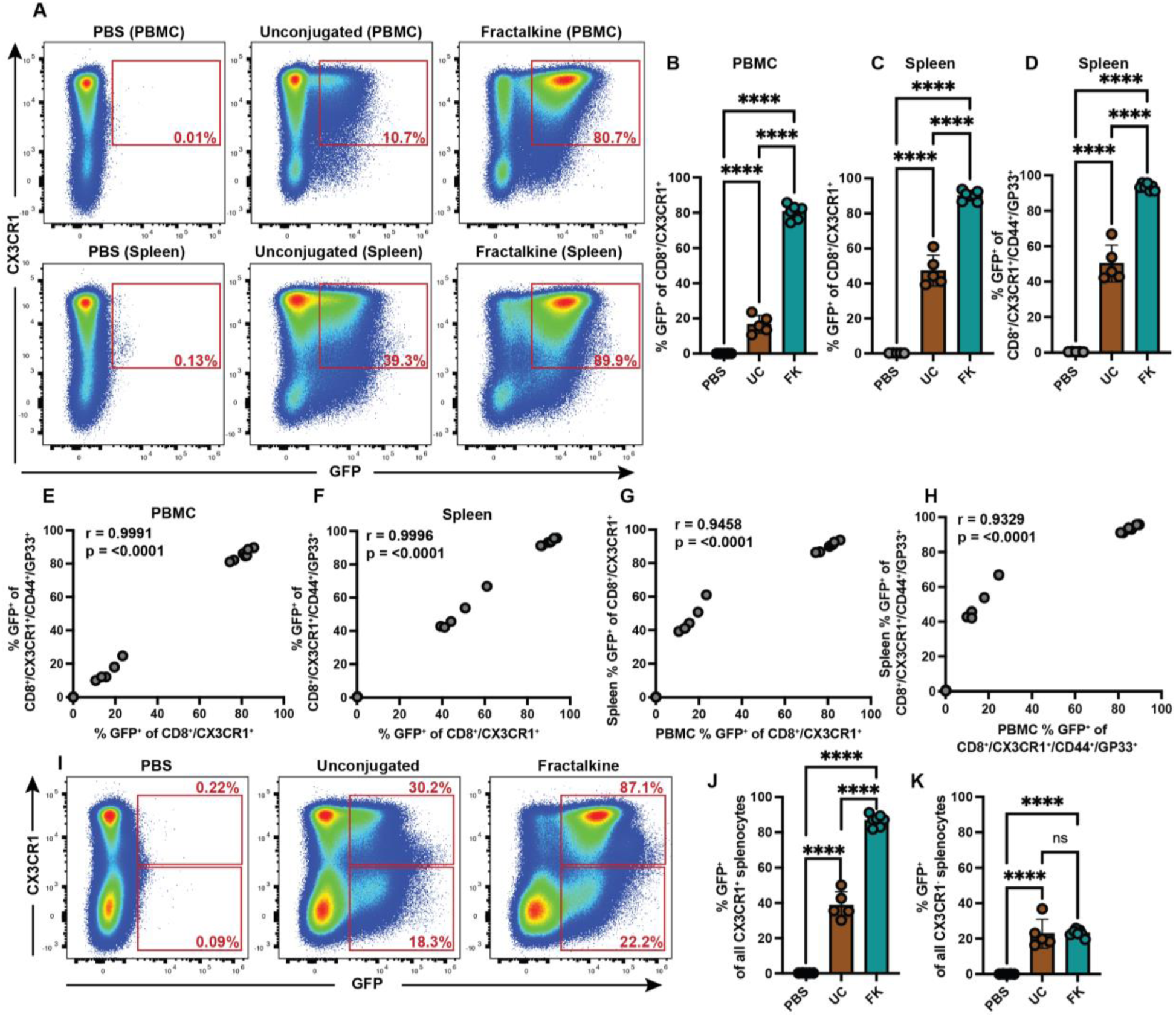
Mouse fractalkine-conjugated LNP target CX3CR1^+^ CD8^+^ T cells in vivo: (A) GFP expression in CX3CR1^+^ CD8^+^ T cells from peripheral blood (top panel) and spleen (bottom panel) 24 hours after PBS or LNP injection (10μg dosage, 1:1.5 μg mRNA/μg fractalkine ratio, IV delivery) with results from one mouse per group shown. Mice were infected with LCMV Armstrong for 8 days prior to treatment. (B) GFP expression in CX3CR1^+^ CD8^+^ T cells from peripheral blood and (C) spleen as shown in (A) and (D) CX3CR1^+^/GP33^+^/CD44^+^/CD8^+^ T cells from spleen 24 hours after PBS, unconjugated GFP LNP or murine fractalkine-conjugated GFP LNP injection (10μg dosage, 1:1.5 μg mRNA/μg fractalkine ratio, IV delivery) for all mice. Mice were infected with LCMV Armstrong for 8 days prior to treatment. (E) Correlation of the percentage of GFP^+^ CX3CR1^+^/GP33^+^/CD44^+^/CD8^+^ T cells and GFP^+^ CD8^+^ T cells from peripheral blood and (F) spleen 24 hours after treatment. (G) Correlation of the percentage of GFP^+^ CD8^+^ T cells from peripheral blood and spleen 24 hours after treatment. (H) Correlation of the percentage of GFP^+^ CX3CR1^+^/GP33^+^/CD44^+^/CD8^+^ T cells from peripheral blood and spleen 24 hours after treatment. (I) GFP expression in CX3CR1^+^ and CX3CR1^-^ splenocytes 24 hours after PBS or LNP injection (10μg dosage, 1:1.5 μg mRNA/μg fractalkine ratio, IV delivery) with results from one mouse per group shown. Values calculated from CX3CR1^+^ and CX3CR1^-^ cells, but flow plots shown for all splenocytes. (J) Expression of GFP in CX3CR1^+^ and (K) CX3CR1-total splenocytes from (I) calculated for all mice included in study. For all statistical analyses, *p < 0.05, **p < 0.01, ***p < 0.001, ****p < 0.0001; ns, not significant.

**Fig. S7.**
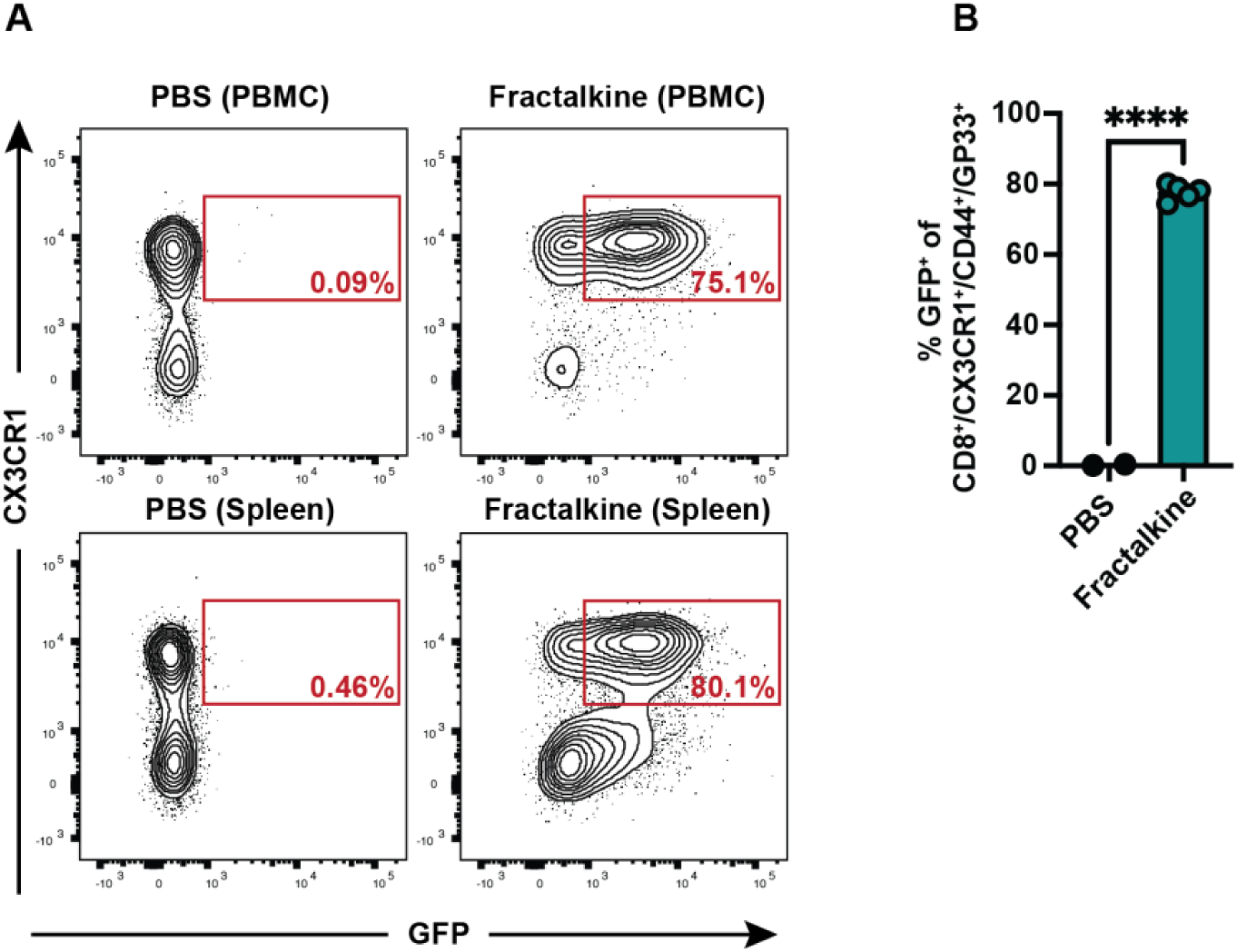
Mouse fractalkine-conjugated LNP target CX3CR1^+^ CD8^+^ T cells after resolution of LCMV Armstrong infection: (A) GFP expression in CX3CR1^+^/GP33^+^/CD44^+^/CD8^+^ T cells from PBMC (top panel) or spleen (bottom panel) 24 hours after injection of either PBS or mouse fractalkine-conjugated GFP LNP (1:0.75 μg mRNA/μg fractalkine ratio, 10μg dosage, IV delivery) with one mouse per group shown. Mice were infected with LCMV Armstrong for 33 days prior to treatment. (B) GFP expression in CX3CR1^+^/GP33^+^/CD44^+^/CD8^+^ T cells from spleen from (A) for all mice included in study. For statistical analysis *p < 0.05, **p < 0.01, ***p < 0.001, ****p < 0.0001; ns, not significant.

**Fig. S8.**
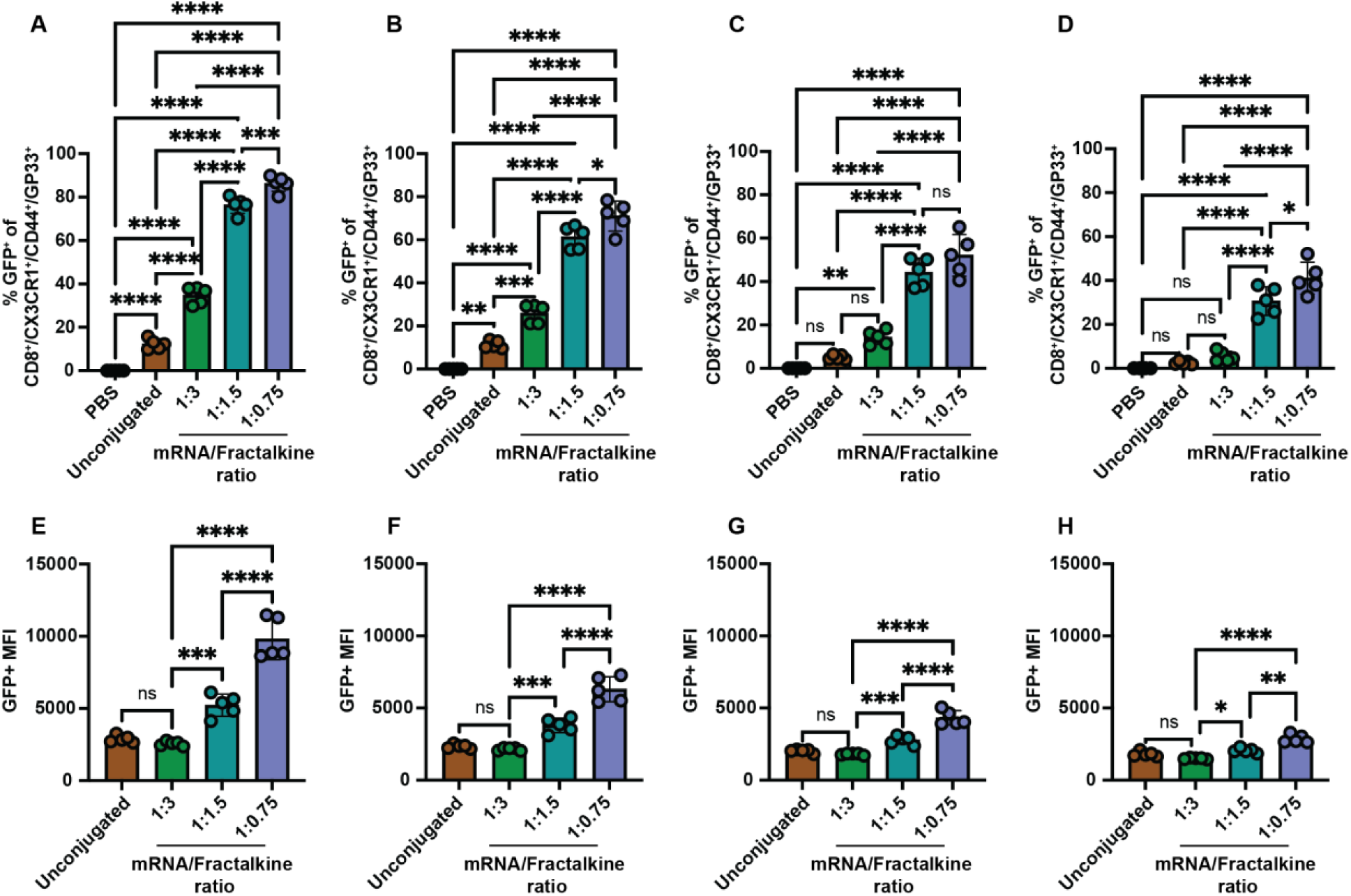
Low fractalkine/mRNA conjugation ratios yield increased GFP expression in CD8^+^ T cells: Percentage of peripheral blood CX3CR1^+^/GP33^+^/CD44^+^/CD8^+^ T cells expressing GFP at (A) 24, (B) 48, (C) 96, and (D) 144 hours after injection of PBS, unconjugated GFP LNP or mouse fractalkine-conjugated GFP LNP at one of three mRNA-to-fractalkine ratios, namely 1:3, 1:1.5 and 1:0.75 μg mRNA/μg fractalkine at a 10μg dosage, IV delivery, for all mice included in study. Mice were infected with LCMV Armstrong for 8 days prior to treatment. Median fluorescence intensity for GFP^+^/CX3CR1^+^/GP33^+^/CD44^+^/CD8^+^ T cells from (A-D) at (E) 24, (F) 48, (G) 96, and (H) 144 hours after injection of unconjugated GFP LNP or mouse fractalkine-conjugated GFP LNP for all mice included in study. For all statistical analyses, *p < 0.05, **p < 0.01, ***p < 0.001, ****p < 0.0001; ns, not significant.

**Fig. S9.**
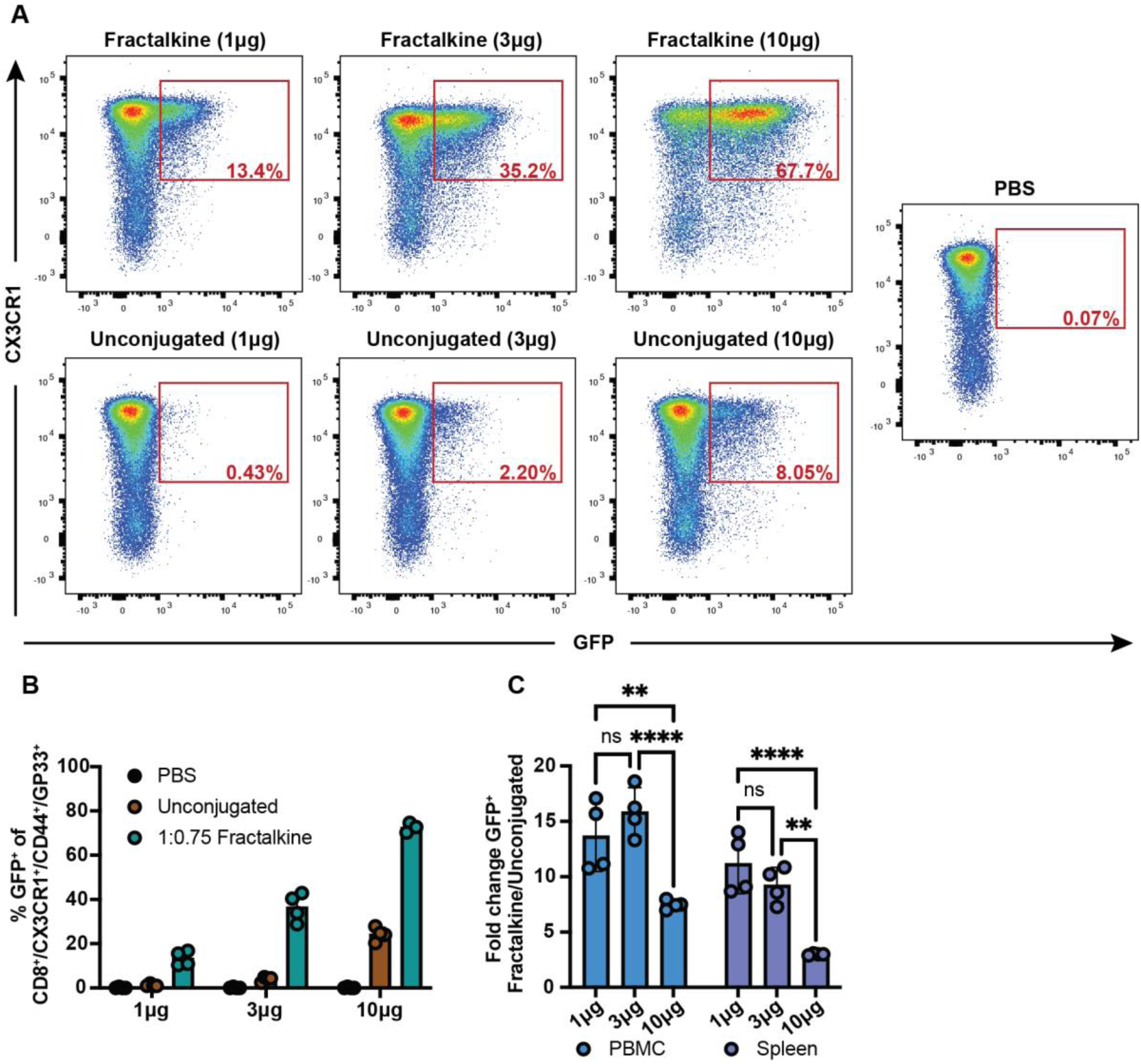
Dose dependent uptake of fractalkine-conjugated mRNA-LNP in vivo: (A) GFP expression in CX3CR1^+^/GP33^+^/CD44^+^/CD8^+^ T cells from PBMC 24 hours after injection of PBS, unconjugated GFP LNP or mouse fractalkine-conjugated GFP LNP (1:0.75 μg mRNA/μg fractalkine ratio, IV delivery) at dosages of 1μg, 3μg, or 10μg with one mouse per group shown. Mice were infected with LCMV Armstrong for eight days prior to treatment. (B) GFP expression in CX3CR1^+^/GP33^+^/CD44^+^/CD8^+^ T cells from spleen 24 hours after injection of PBS, unconjugated GFP LNP or murine fractalkine-conjugated GFP LNP (1:0.75 μg mRNA/μg fractalkine ratio, IV delivery) at dosages of 1μg, 3μg, or 10μg for all mice included in study. (C) The fold change in %GFP^+^ CX3CR1^+^/GP33^+^/CD44^+^/CD8^+^ T cells following treatment with fractalkine-conjugated versus unconjugated LNP at the indicated doses in either PBMCs or splenocytes. Individual values of %GFP^+^ CX3CR1^+^/GP33^+^/CD44^+^/CD8^+^ T cells from the fractalkine-conjugated LNP groups were divided by the average %GFP^+^ CX3CR1^+^/GP33^+^/CD44^+^/CD8^+^ T cells from the corresponding unconjugated LNP group.

**Fig. S10.**
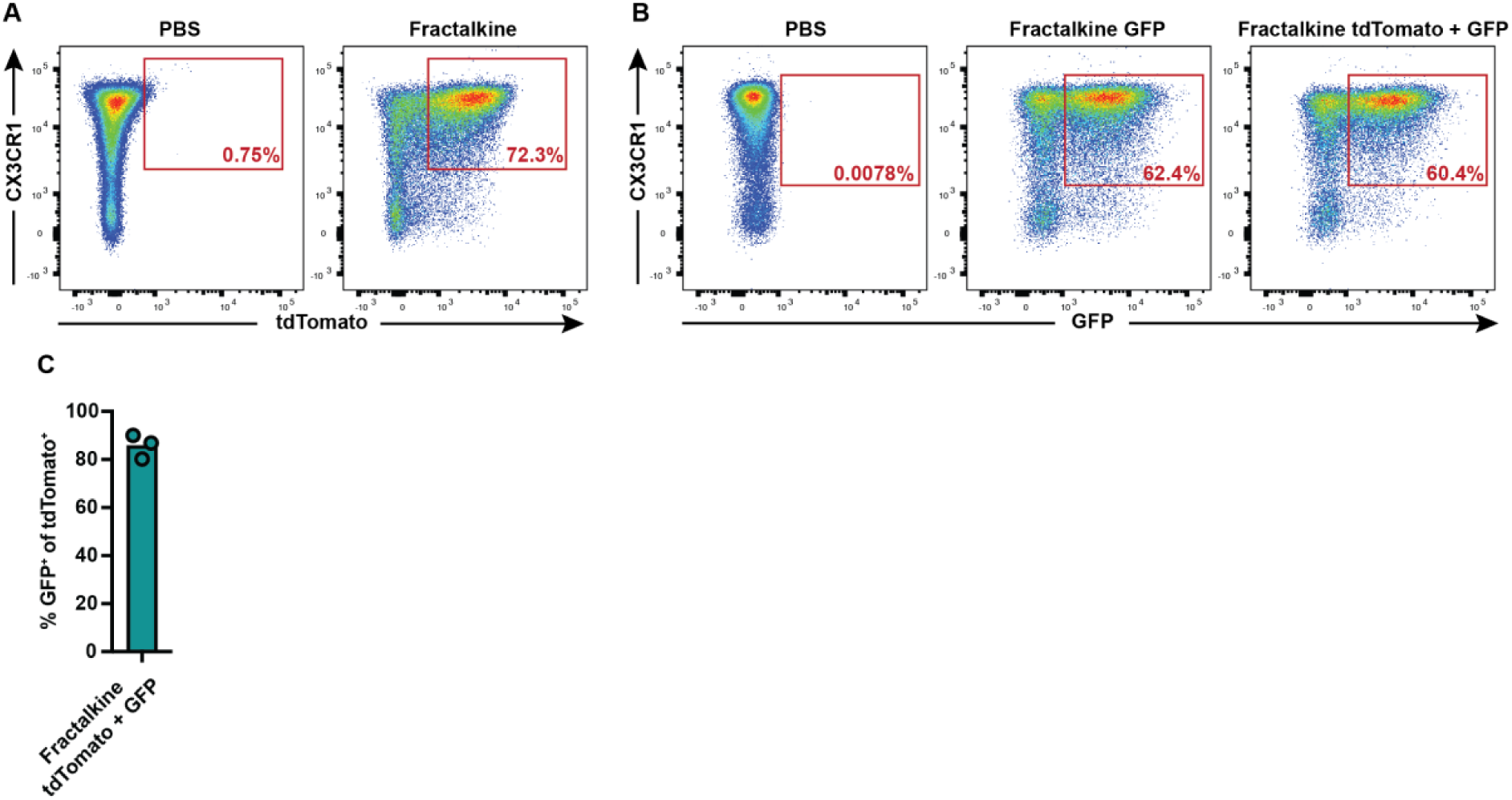
Fractalkine-conjugated mRNA-LNP sequentially target CX3CR1^+^ CD8^+^ T cells in vivo: (A) tdTomato expression in CX3CR1^+^/GP33^+^/CD44^+^/CD8^+^ T cells from PBMC 24 hours after injection of PBS or mouse fractalkine-conjugated tdTomato LNP (1:0.75 μg mRNA/μg fractalkine ratio, 10μg dosage, IV delivery) with one mouse per group shown. Mice were infected with LCMV Armstrong for 7 days prior to treatment. (B) GFP expression in CX3CR1^+^/GP33^+^/CD44^+^/CD8^+^ T cells from PBMC 24 hours after retro-orbital injection of PBS or mouse fractalkine-conjugated GFP LNP (1:0.75 μg mRNA/μg fractalkine ratio, 10μg dosage, IV delivery) with one mouse per group shown. Mice were infected with LCMV Armstrong for 8 days prior to treatment and were previously treated with PBS, mouse fractalkine-conjugated tdTomato LNP or nothing one day prior. (C) Percentage of peripheral blood tdTomato^+^/CX3CR1^+^/GP33^+^/CD44^+^/CD8^+^ T cells also expressing GFP on day 9 for all mice included in study.

**Fig. S11.**
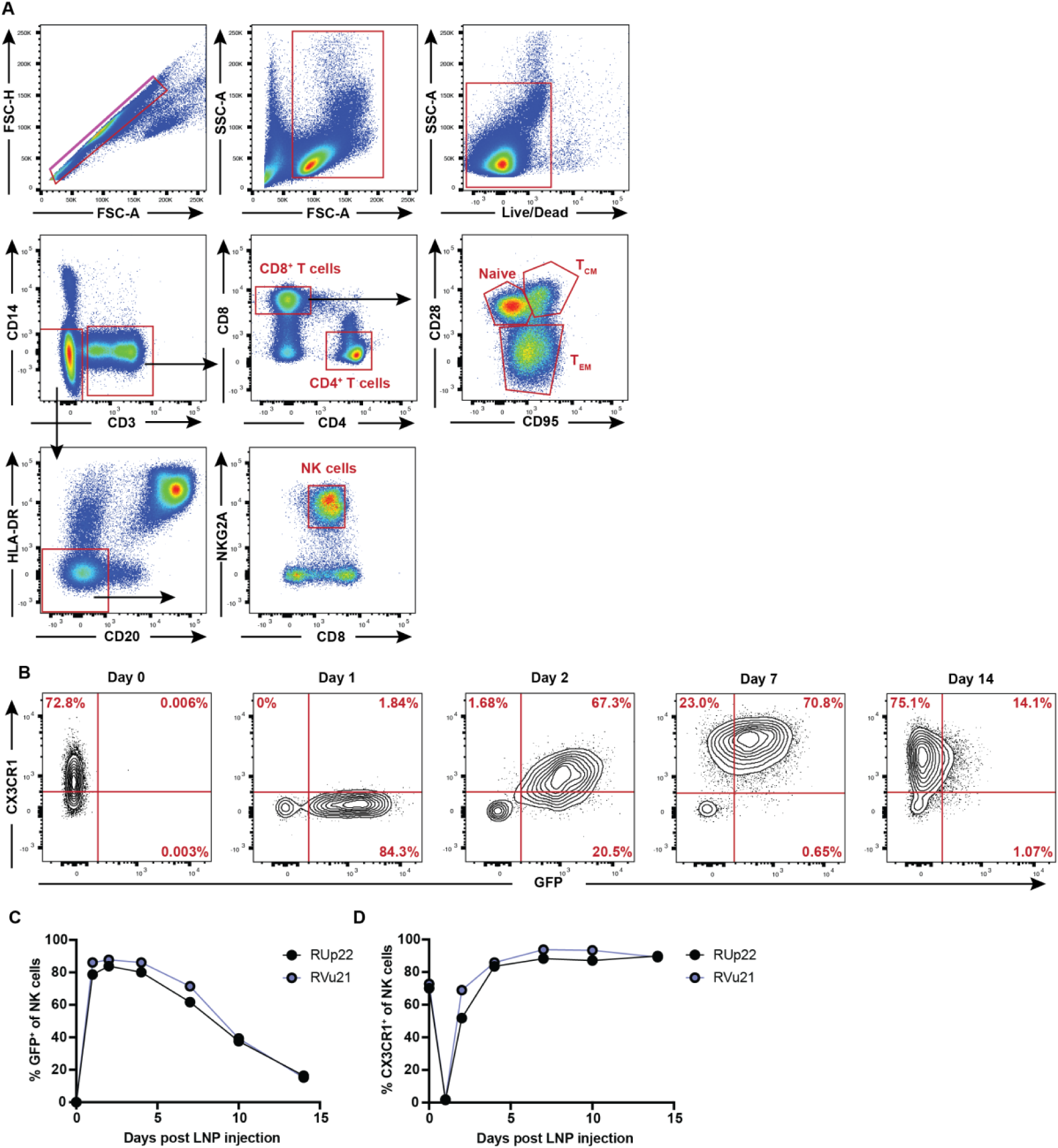
Fractalkine-conjugated mRNA-LNP target NHP CX3CR1^+^ NK cells in vivo: (A) Flow cytometry gating strategy for NHP CD8^+^ T cell and NK cell analysis. CD28 by CD95 gate also shown in Fig. 4E. (B) GFP expression in peripheral blood NK cells either at baseline (Day 0) or after LNP injection (Days 1-14) (0.7mg/kg dosage, 1:2 μg mRNA/μg fractalkine ratio, IV delivery), results from one animal shown. (C) Percentage of peripheral blood NK cells expressing GFP or (D) CX3CR1 across all timepoints analyzed for each animal.

**Fig. S12.**
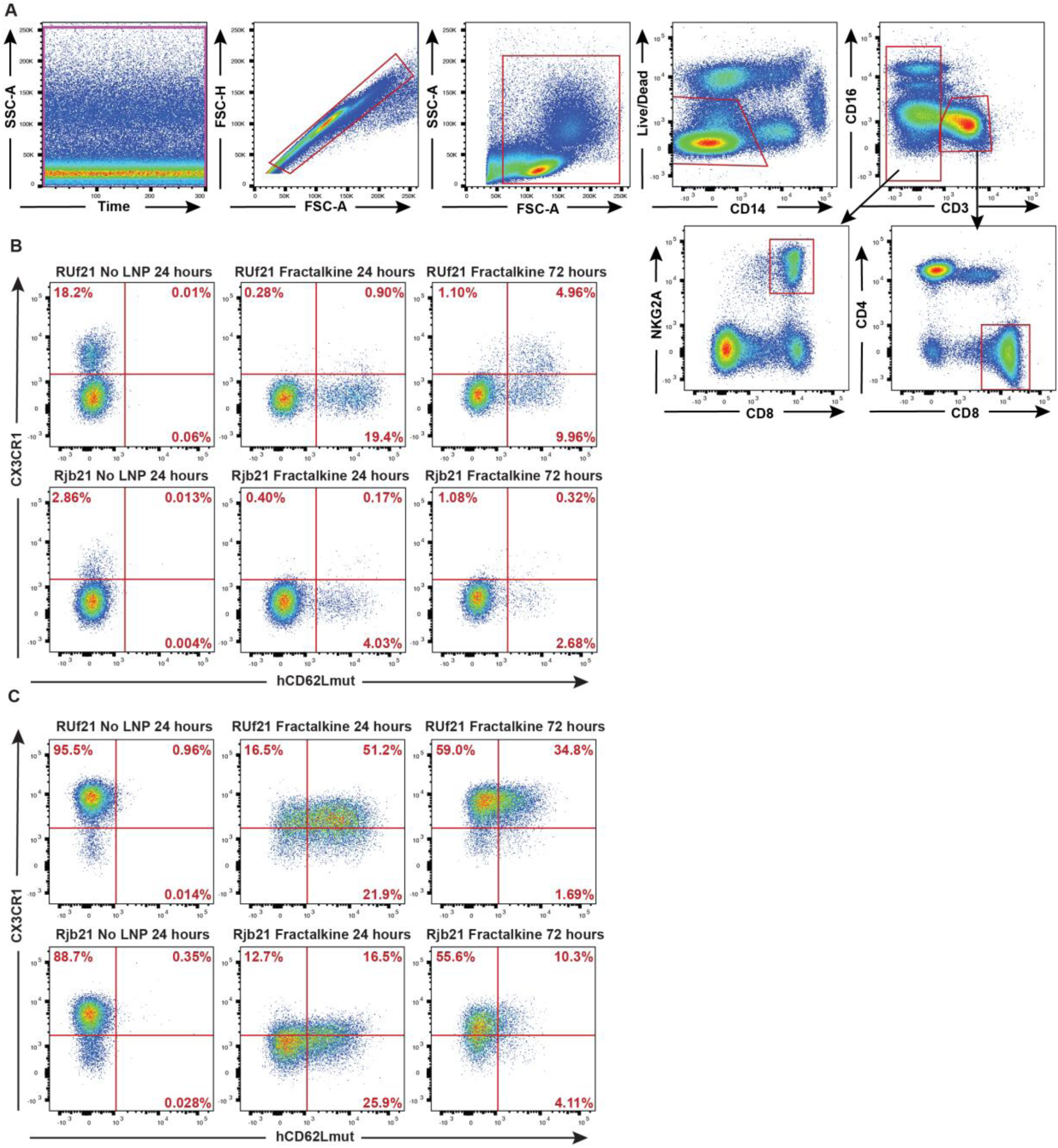
Fractalkine-conjugated human CD62L mRNA-LNP drives hCD62Lmut expression in vitro: (A) Flow cytometry gating strategy for analyzing NHP peripheral blood CD8^+^ T cells and NK cells in vitro. (B) Flow cytometry analysis of purified NHP PBMCs incubated for 24 or 72 hours with or without fractalkine-conjugated human CD62L-encoding mRNA-LNP (0.1µg mRNA/million cells). Expression of CX3CR1 and human CD62L in the (B) CD8^+^ T cells and (C) NK cells from the two NHPs included in the CD62L in vivo study, RUf21 and Rjb21.

**Fig. S13.**
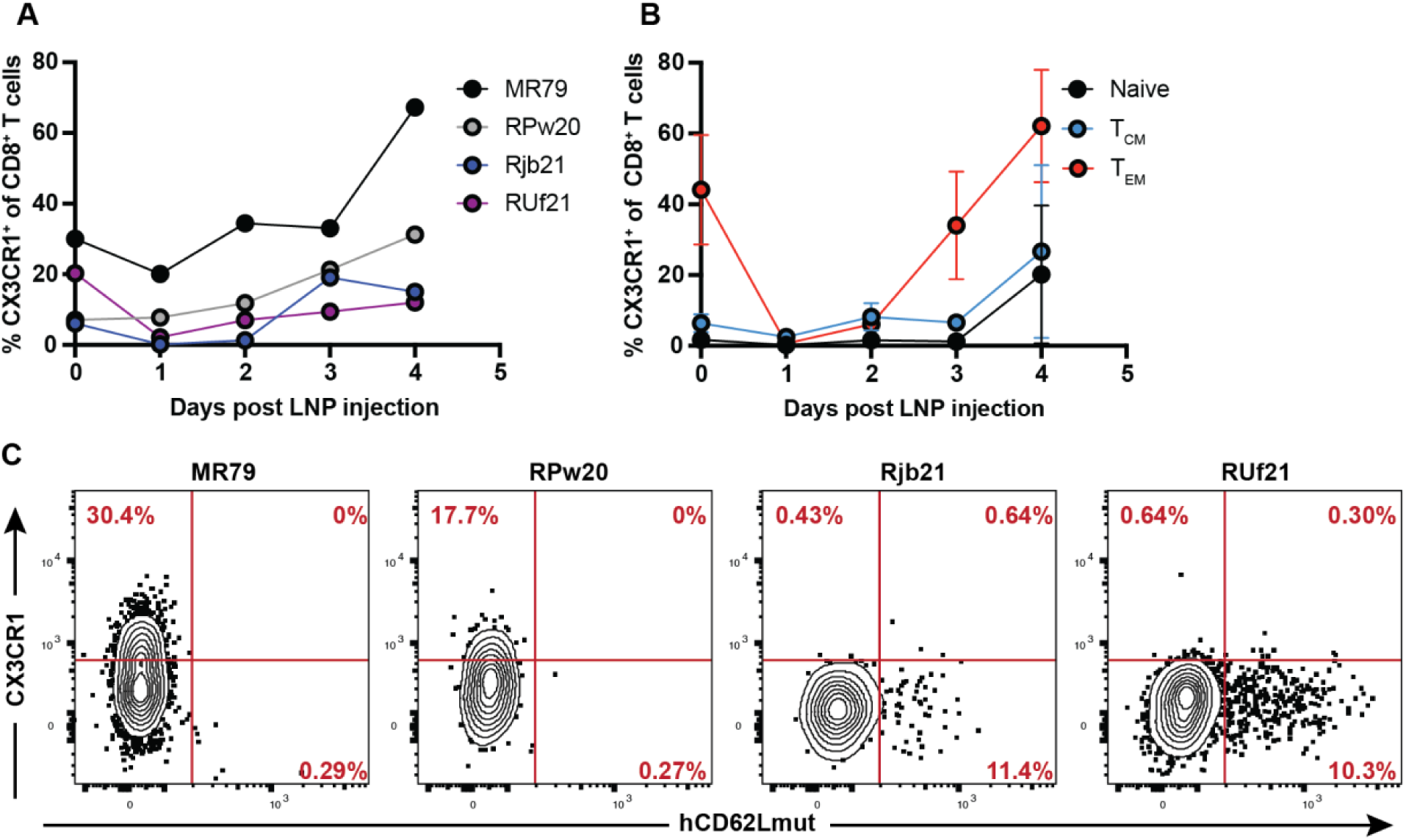
Fractalkine-conjugated hCD62L mRNA-LNP drives hCD62Lmut expression in CX3CR1^+^ Teff cells in vivo: (A) Percentage of NHP peripheral blood CD8^+^ T cells expressing CX3CR1 across all timepoints analyzed. (B) Percentage of peripheral blood CD8^+^ T cells expressing CX3CR1 in each CD8^+^ T cell memory subset across all timepoints analyzed. (C) hCD62Lmut and CX3CR1 expression in lymph node effector memory (TEM; CD28^-^/CD95^+^) CD8^+^ T cells 24 hours after LNP injection.

**Fig. S14.**
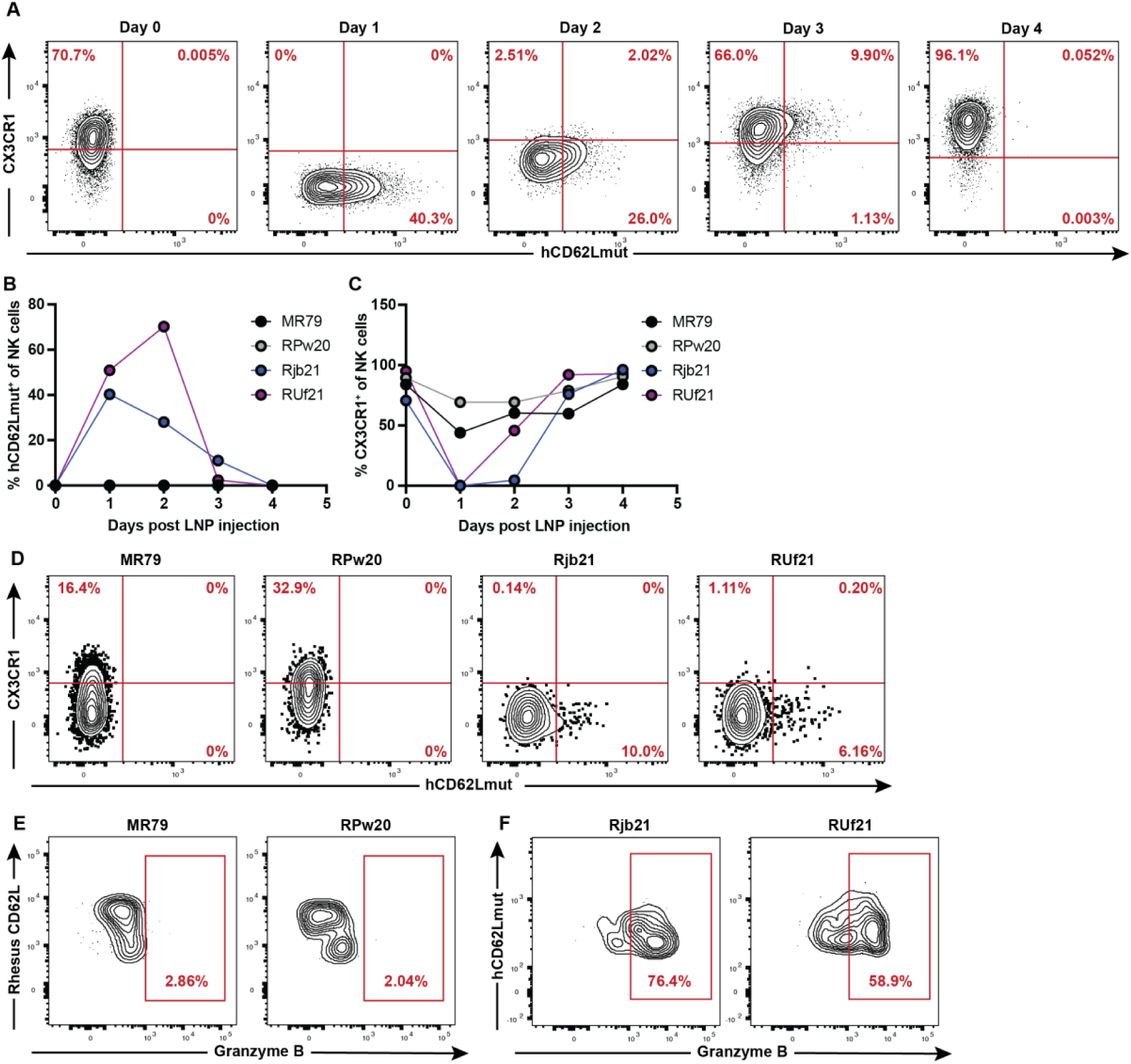
Fractalkine-conjugated hCD62Lmut mRNA-LNP drives hCD62Lmut expression in CX3CR1^+^ NK cells in vivo: (A) hCD62Lmut expression in NHP peripheral blood NK cells either at baseline (Day 0) or after LNP injection (Days 1-4) (0.5mg/kg dosage, 1:2 μg mRNA/μg fractalkine ratio, IV delivery), results from one animal shown. (B) Percentage of peripheral blood NK cells expressing hCD62Lmut or (C) CX3CR1 across all timepoints analyzed. (D) hCD62Lmut and CX3CR1 expression in lymph node NK cells 24 hours after LNP injection. (E) Granzyme B expression in rhesus CD62L^+^ or (F) hCD62Lmut^+^ NK cells from lymph node 24 hours post LNP injection.

